# Nutrient response strategies drive coastal range shifts of phytoplankton taxa

**DOI:** 10.64898/2026.01.13.699386

**Authors:** Arianna I. Krinos, Margaret Mars Brisbin, Amy Costa, Sara K. Shapiro, Michael J. Follows, Mara A. Freilich, Harriet Alexander

## Abstract

Coastal phytoplankton blooms are important drivers of regional and global marine primary production. Recently documented increases in coastal phytoplankton blooms in the first quarter of the 21st century may be a consequence in part of changing environmental conditions and have important ecological implications. Here, we explore coastal phytoplankton dynamics in Cape Cod Bay, MA, USA. We use a 20-year phytoplankton ecology dataset to identify potential drivers of the increasing prevalence of two regionally-rare phytoplankton taxa: a coccolithophore thought to thrive in the global open ocean, and a dinoflagellate genus with potentially toxic members. Using metatranscriptomics, we show that these minor phytoplankton taxa leverage unique strategies to gain a competitive advantage under nutrient limitation compared to traditionally dominant taxa and compared to a diatom taxon that became modestly more abundant over the study period. Our results highlight the ecological dynamics arising from long-term shifts in temperature and nutrient status in coastal ecosystems.

## 1 Introduction

Anthropogenic changes in nutrients and warming water temperatures are shifting the timing, abundance (Thompson et al., 2009), and composition (Camarena-Gómez et al., 2018; Barth et al., 2020) of phytoplankton in coastal environments (Dai et al., 2023; James et al., 2022; Fernández-Alías et al., 2022). Phytoplankton drive the ecology and biogeochemistry of coastal ecosystems and shifts in phytoplankton ecology have been observed to propagate up the food chain with implications for coastal fisheries (Townsend, 1991; Pershing et al., 2021). Changes in dominant phytoplankton community members may alter biogeochemistry through changing carbon cycling and export patterns (Landa et al., 2016; Smith Jr et al., 2021), contributing to increases in seasonal bottom oxygen depletion (Scully et al., 2022), or to toxin production (Glibert et al., 2016). While recent trends of increases in phytoplankton bloom frequency (Dai et al., 2023) have been attributed to rising temperature, shifts in nutrient availability due to changing circulation and anthropogenic nutrient supply may be particularly profound in coastal environments (Oczkowski et al., 2018).

Changing nutrient regimes modify not only the abundance of phytoplankton but also their community composition. Climate change has resulted in the emergence of previously rare or undocumented phytoplankton groups in coastal environments (Scully et al., 2022). While changing phytoplankton taxonomy suggests altered community function, causal mechanisms for shifts in community composition and definitive evidence of altered function are now considerably more accessible with molecular tools (Mock et al., 2016). Metatranscriptomes can detect low-abundance populations and probe the putative metabolic activity of these minor taxa (Zampieri et al., 2023). This is especially important because routine phytoplankton count data can fail to identify growing, metabolically-active, and diverse populations. Phytoplankton counts may miss rare taxa and lack in taxonomic resolution (for example, cryptic species (Smayda, 2011)), in addition to missing small, difficult to visualize or distinguish cells, and omitting metabolic information. Monitoring metabolism in situ via metatranscriptomics provides a community context for the expression of genes involved in environmental response (Moran et al., 2013). Understanding phytoplankton metabolism and how it responds to local trends can help identify traits that differentiate biogeochemically-relevant phytoplankton taxa and predict how these groups may continue to change in abundance in the future.

This study, centered in coastal Cape Cod Bay (CCB), presents the first -omic data coupled to a long-term study of the ecology of this changing coastal ecosystem, and has broader implications for changing coastal ecosystems similar to CCB. We document the shift towards two formerly rare phytoplankton taxa in ecologically-important coastal waters via metatranscriptomic time series collected to contextualize long-term phytoplankton abundance data from counts, and compare the metabolism of those formerly rare taxa to that of other groups. We focus on coccolithophores and dinoflagellates in the genus *Prorocentrum*, both of which are groups that have historically low abundance in CCB, yet with key ecosystem roles: coccolithophores are calcifying phytoplankton contributing to both organic and inorganic carbon cycling (Paasche, 2001), and dinoflagellates including *Prorocentrum* may be invasive toxin-producers (Telesh, Schubert, and Skarlato, 2016; Hallegraeff, 2003; Faust and Gulledge, 2002). The cosmopolitan coccolithophore *Gephyrocapsa huxleyi* has been observed on the continental shelf beyond CCB, and some of the cultured strains deposited in culture collections have been obtained from this region (Fielding, 2013; Marschall, 1985).

Nutrient depletion in summer surface waters—driven by spring-bloom phytoplankton (e.g., *Phaeocystis* in bloom years (Smith Jr et al., 2021))—could promote the growth of specific phytoplankton groups adapted to the warm, low-nutrient conditions of the North Atlantic subtropical gyre, including haptophytes like coccolithophores that are suited to oligotrophic conditions. Alternatively, small mixotrophic generalists like the dinoflagellate *Prorocentrum* may outcompete other taxa in the warm, nutrient-depleted summer waters due to their flexibility in using nutrients that may be inaccessible to other phytoplankton groups. Studying coccolithophore and dinoflagellate growth in response to warming and more nutrient-limited conditions in CCB informs how warm temperature-acclimated, nutrient-stressed populations behave metabolically.

Our monthly metatranscriptomic dataset also circumvents the challenge that low overall abundances make taxa challenging to accurately enumerate via microscopy. This is especially true for coccolithophores like *G. huxleyi*, which have relatively low average cell diameter, and multiple morphologies due to calcification state and ploidy, and hence may not always be possible to visually identify and differentiate from other taxa. Metatranscriptomic time series coupled to other long-term ecological data illuminate potential mechanisms for observed shifts in phytoplankton community composition in coastal environments. Community gene expression data provide possible metabolic mechanisms that underlie changing biodiversity as a consequence of climate change and other anthropogenic impacts on sensitive coastal ecosystems.

## 2 Materials and Methods

### 2.1 Field sampling by Center for Coastal Studies and MWRA

Since 1992, long-term monitoring at field sites in Cape Cod Bay has been funded by the Massachusetts Water Resources Authority (MWRA) monitoring program (Hunt et al., 2010), which has varied in spatial extent during the time period from 1992 to 2023. Since 1999, the Massachusetts Water Resources Authority (MWRA) has collected phytoplankton and zooplankton count and nutrient and physical data from a series of gridded sampling points in CCB. Beginning in 2011, MWRA contracted the Center for Coastal Studies (CCS) in Provincetown, MA to conduct this sampling at 3 of these stations, 2 in CCB, and one in Stellwagen Bank National Marine Sanctuary. These sites are visited monthly to assess water quality and microbial biodiversity in CCB. In this study, we focused on the three stations in CCB where microscopic phytoplankton counts have been collected. The methods for sample collection at the CCB monitoring sites are described in detail in (Hunt et al., 2010) and in the technical reports published for the sampling site (Libby et al., 2006; Yard et al., 2002; Prasse et al., 2004; Costa et al., 2023; Costa, Larson, and Stamieszkin, 2013). In brief, monthly hydrographic data were collected using a conductivity-temperature-depth (CTD) instrument with additional sensors for photosynthetically active radiation (PAR), chlorophyll fluorescence, and dissolved oxygen. Surface water samples were collected for phytoplankton enumeration and were fixed with alkaline Lugol’s solution (“100 g potassium iodide, 50 g iodine, and 50 g sodium acetate each are dissolved incrementally in distilled water to a final volume of 1 L” per (Costa et al., 2023)). A vertical net tow over the upper 19 meters was collected for formalin-fixed zooplankton enumeration. Chlorophyll *a* was also measured at each site to calibrate sensor-based fluorescence measurements. After vacuum-filtering water samples onto glass fiber filters (47-mm GF/F filters), the filters were frozen and chlorophyll was measured with a fluorometer (Turner Designs) (Yentsch and Menzel, 1963; Arar and Collins, 1997).

Dissolved inorganic nutrients were measured by filtering water samples through 0.4-*µ*m membrane filters and freezing the sample. Colorimetric analysis was used to determine concentrations of nitrate, nitrite, and ortho-phosphate according to the methods described in the quality assurance reports published by the MWRA and the Center for Coastal Studies (Costa, Larson, and Stamieszkin, 2013). Phytoplankton were enumerated from 800 mL water samples after settling to 50 mL and vacuum aspiration, resulting in a 1 mL aliquot counted using an Olympus BH-2 compound microscope with phase contrast (Hunt et al., 2010; Costa, Larson, and Stamieszkin, 2013). Following visual identification, total concentration of phytoplankton in the sample was estimated based on original volume converted to cells per liter. Zooplankton abundance was estimated in organisms per cubic meter based on subsampling at least 250 organisms to identify with a Leica L2 stereomicroscope and performing a calculation based on the flow meter change during the tow and the area of the net opening (Costa, Larson, and Stamieszkin, 2013).

### 2.2 Sampling for metatranscriptomics

We collected water samples and sequenced metatranscriptomes from the MWRA station in Stellwagen Bank for a period of three field seasons, with samples collected monthly between February 2021 and July 2023, and sequenced metatranscriptomes from the two remaining stations in July 2021 and July 2023 (Table 4). A sample of water of approximately 20 liters was collected at approximately 1 meter depth via repeated water collection with a Van Dorn sampler. The water sample was stored in a cleaned polycarbonate carboy covered in black plastic film to avoid light exposure prior to filtering. On February sampling dates, ice packs were used to minimize the temperature change of the carboys. The water samples were taken back to the laboratory for filtration using a peristaltic pump. After cycling one liter of water to rinse and prime the filtering setup, approximately 5 liters of water per each of three replicates were filtered (depending on the density of the sample, which influenced the ease of filtration; total volume filtered was documented). The water was pumped through two sequential mixed cellulose ester (MCE) filters, the first of which had a 47 millimeter 0.8 *µ*m pore size and the second of which had a 0.2 *µ*m pore size (47 millimeter diameter; MilliporeSigma). The filters, collected in triplicate, were sliced into halves with a sterilized razor blade. The two halves were stored in separate plastic cryogenic storage vials prior to being flash frozen in liquid nitrogen and subsequently stored in a *−*80*°*C freezer until further analysis.

### 2.3 RNA extraction and sequencing

RNA was extracted from the stored and frozen samples using an initial bead beating step with BashingBead lysis tubes (Zymo Corporation, CA, USA) containing 0.1 and 0.5 nm chemically inert dry beads, to which each filter and the initial lysis buffer (beta-mercaptoethanol (*β*ME) and RLT Master Mix from the RNeasy extraction kit, Qiagen) was added. The tubes were incubated at 65*°*C for ten minutes prior to vortexing at high speed for 5 minutes using a vortex tube adapter. The lysis tubes were then centrifuged for 1 minute at 4,000 *×* g (rcf) and the supernatant was used as input to the RNeasy kit protocol (Qiagen) performed according to the manufacturer’s instructions, including the DNase digestion step. A NEBNext Ultra II Directional RNA Library Prep Kit for Illumina (New England BioLabs, MA, USA) was used for library preparation according to manufacturer protocol, including adding barcode indices for Illumina sequencing on a single sequencing lane. RNA was poly-A selected to filter for eukaryote mRNAs using the poly-A mRNA magnetic isolation module. Approximately 45 samples per lane were sequenced on an Illumina NovaSeq S4 run at the Columbia Genome Center with paired-end 100 basepair reads. All monthly samples collected at station F29 were processed and sequenced in triplicate, and the additional samples in July 2021 and July 2023 from stations F01 and F02 further south in Cape Cod Bay were targeted and sequenced in triplicate based on review of the microscopic count results for coccolithophores (Table 4).

### 2.4 Metatranscriptome assembly and post-processing

Short reads from all metatranscriptomic samples (Table 4) were pre-processed, assembled and annotated with the *euk*rhythmic pipeline (Krinos et al., 2023). Reads were trimmed using trimmomatic version 0.39 in paired-end mode (Bolger, Lohse, and Usadel, 2014) (parameters: ILLUMINACLIP: <STANDARD-adapter-file>:2:30:7 LEADING:2 TRAILING:2 SLIDINGWINDOW:4:2 MINLEN: 50, using a standard list of Illumina adapters as available for download with the *euk*rhythmic pipeline (Krinos et al., 2023)), and quality was checked using FastQC (Andrews et al., 2012) with default parameters. Assembly was performed using MEGAHIT version 1.2.9 (Li et al., 2015) (parameters: minimum *k*-mer value of 20 and maximum *k*-mer value of 110, -m 0.9 -t 8) and rnaSPAdes version 3.15.5 (Bushmanova et al., 2019) (with default parameters other than increasing maximum threads (-t) to up to 20 and maximum memory (-m) to up to 1 terabyte). Assembled contigs were co-clustered at a coverage cutoff of the shorter sequence of 95% using mmseqs2 (Steinegger and Söding, 2017) version 15.6f452 (using linclust, parameters: --min-seq-id 1.00 --cov-mode 1 -c 0.95 --split-memory-limit 120G --remove-tmp-files). Final assembled and clustered contigs were taxonomically annotated using EUKulele version 2.0.7 (Krinos et al., 2021) using a database containing the Marine Microbial Eukaryote Transcriptome Sequencing Project transcriptomes (Keeling et al., 2014; Johnson, Alexander, and Brown, 2019) and the MarRef database (Klemetsen et al., 2018). Open reading frames were predicted on the assembled contigs using TransDecoder version 5.5.0 (Haas and Papanicolaou, 2021) (parameters: -m 100) and functional annotations were performed using eggNOG-mapper version 1.0-190-g11acf21 (Huerta-Cepas et al., 2017) (completed in 30 separate subfiles for initial hits to orthologs and then combined using --annotate_hits_table; parameters: -m diamond --override --no_annot --no_file_comments --cpu 18 for initial hits command and then --no_file_comments --cpu 18 for annotation of the hits table). The mapping rate of raw read sequences was estimated using the Salmon mapping tool version 1.4.0 (Patro et al., 2017) (parameters: -p 20 --validateMappings for index creation and -l A -p 20 –validateMappings for the quantification step).

### 2.5 Sequence data analysis

Assemblies were subsetted based on the taxonomic group of interest as follows, via the taxonomic annotations from EUKulele (Krinos et al., 2021). Coccolithophores were defined as transcripts annotated as belonging to family Noerlaerhabdaceae, Leptocylindrales diatoms were defined as transcripts belonging to genus Leptocylindrus, Thalassiosirales diatoms were defined as transcripts belonging to family Thalassiosiraceae, cryptophytes were defined as transcripts belonging to order Cryptomonadales, and *Prorocentrum*/Prorocentrales dinoflagellates were defined as transcripts belonging to order Prorocentrales. After transcripts were subsetted based on taxonomic group, functional annotations were performed on the predicted peptide sequences from each group of transcripts using KofamScan (Aramaki et al., 2020). Abundance was assigned to each transcript in each sample using estimated transcripts per million (TPM) abundances via Salmon (Patro et al., 2017), and then normalized via a *Z*-score metric calculated based on mean and standard deviation of the abundance of the individual taxonomic group. Differential expression analysis was performed using DESeq2 (Love, Huber, and Anders, 2014).

### 2.6 Chlorophyll trend analysis

Regional trends in chlorophyll were quantified using daily MODIS Aqua L3 satellite product at 4.63 km resolution. Chlorophyll is computed using the OCI algorithm (Hu, Lee, and Franz, 2012). Trends are computed as a linear fit at each pixel in the timeseries with significance assessed using a t-test.

### 2.7 Nutrient and temperature data analysis and correlations

Nutrient and temperature data were collected at multiple depths; depths below 5 meters were considered “surface” and averaged for the purposes of Spearman correlation calculation, which was performed in R version 4.3.1. For nitrate and nitrite, prior to October 2010, nitrogen molar concentrations from nitrite and nitrate were measured separately, and after this timepoint, combined nitrite and nitrate-derived nitrogen was reported. As a result, when nitrite and nitrate were measured separately, the two values were summed at each depth of measurement prior to averaging.

## 3 Results

### 3.1 Minor taxa emerge in the context of increasing coastal chlorophyll

The Gulf of Maine is a well-studied coastal ecosystem that has provided ecosystem services to the local population for centuries via a productive fishery (Pershing et al., 2015). Recently, this region has experienced significant sea surface temperature warming that has led to ecological shifts, and may result in a future fishery collapse (Pershing et al., 2015; Pershing et al., 2021). In an average year, Cape Cod Bay (CCB) follows the Gulf of Maine and North Atlantic ecological pattern of spring phytoplankton bloom, followed by grazer predation, followed by summer seasonal nutrient limitation (Townsend, 1991).

Since 2010, summer (June 21 through September 20) chlorophyll has increased modestly in CCB (Figure 1); this trend was significant at the two CCB stations (F01: *R*^2^ = 0.10, *p* = 0.021, slope of chlorophyll in *µ*g L*^−^*^1^ vs. months since 2010 = 0.012; F02: *R*^2^ = 0.11, *p* = 0.016, slope of chlorophyll in *µ*g L*^−^*^1^ vs. months since 2010 = 0.005), but not at the Stellwagen Bank station (F29: *p*=0.10). A marginally significant trend was identified for chlorophyll across all months rather than just for the summer months at station F29 in Stellwagen Bank (F29: *R*^2^ = 0.026, *p* = 0.043, slope of chlorophyll in *µ*g L*^−^*^1^ vs. months since 2010 = 0.007). When correlations were calculated separately only for measurements taken in the month of July, all three stations showed significant correlations of chlorophyll with months since 2010 (F01: *R*^2^ = 0.39, *p* = 0.13, slope = 0.007; F02: *R*^2^ = 0.49, *p* = 0.005, slope = 0.005; F29: *R*^2^ = 0.58, *p* = 0.001, slope = 0.007). In the satellite data, all locations in the grid of the latitude and longitude bounding box around the three stations we sampled in CCB had a significant increasing trend; the range of least-squares polynomial fits in this region was an increase in chlorophyll of 0.048 to 0.11 mg m*^−^*^3^ year*^−^*^1^ (mean: 0.070 mg m*^−^*^3^ year*^−^*^1^).

**Figure 1:**
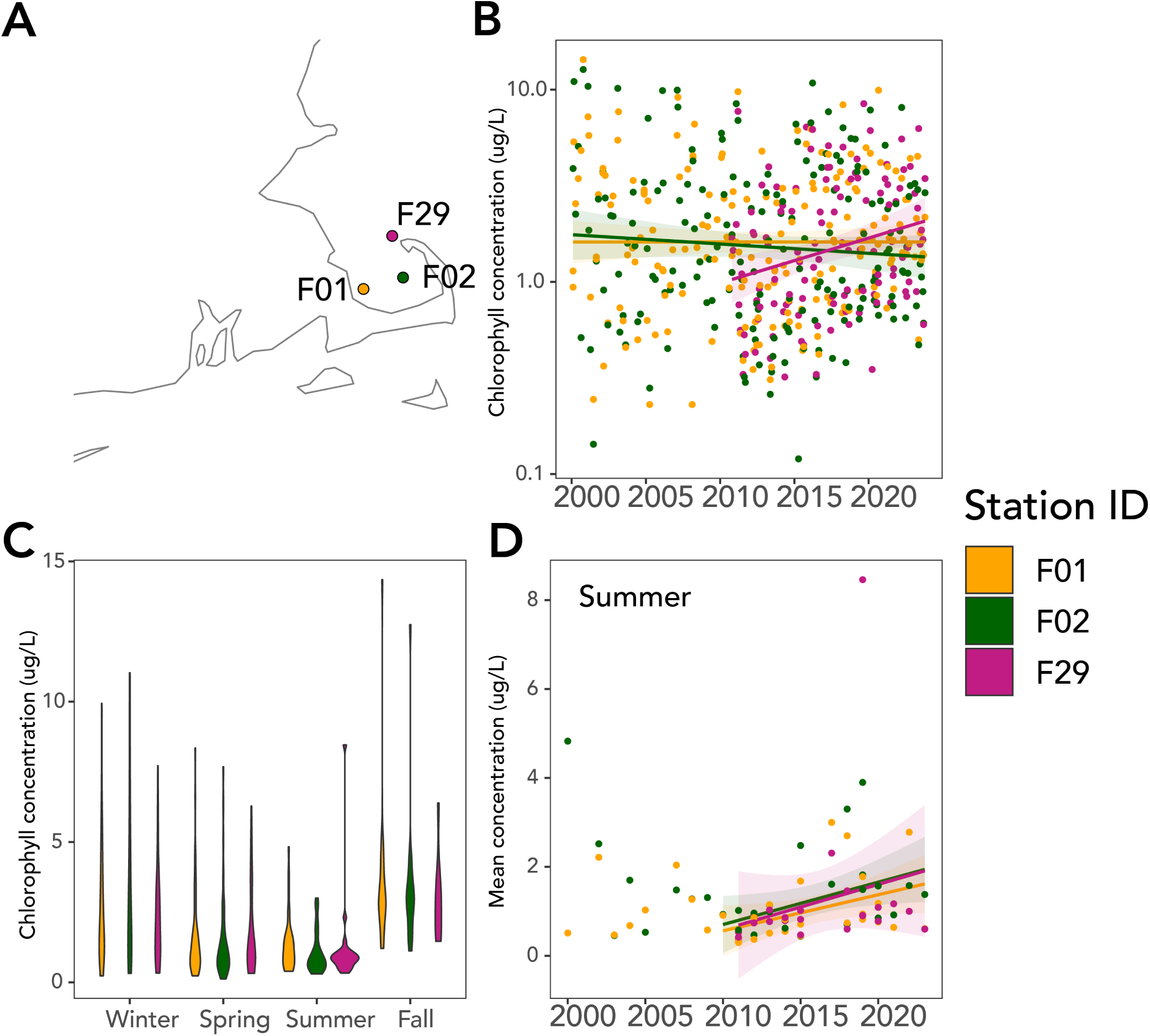
Chlorophyll trends over time in CCB. Chlorophyll trends over time in Cape Cod Bay (CCB). A: Map of sampling sites in CCB colored by station. Tile colors show correlation between time and chlorophyll estimated from satellite data between 2002 and 2025 using the coefficient of the estimated linear fit at each assessed grid cell. Scale is between −0.5 and 0.5; tiles that exceed this range in areas outside of Cape Cod Bay are shaded in gray with white asterisks (*) for values less than −0.5 and orange asterisks (*) for values greater than 0.5. B: Chlorophyll concentration in *µ*g L*^−^*^1^ over the sampling period from 2000-2023 (F01 and F02) and 2010-2023 (F29). C: Violin plots showing distribution of chlorophyll concentrations in each season for each of the stations, illustrating low chlorophyll values in summer. D: Trend in summer-only chlorophyll values in CCB, showing a post-2010 increase in chlorophyll.

In recent years, summer nutrient limitation has become more pronounced in CCB (Figure 2; significance test of post-2010 compared to 2010 and earlier for orthophosphate: *t* = *−*11.8; *p <* 1*e^−^*^15^; nitrate and nitrite: *t* = *−*5.4; *p <* 1*e^−^*^7^), often with large surface phytoplankton blooms and subsequent dissolved oxygen depletion (Scully et al., 2022). Coccolithophores and *Prorocentrum* both increased in summertime abundance in 2010-2023 as compared to 2000-2010 in concert with an overall decrease in inorganic macronutrient (nitrogen and phosphorus) availability, a shift which occurred principally after 2010 (Figure 3). All the highest recorded cell concentrations of coccolithophores in the time series occurred after 2016 (Figure 3; two-sample t-test, alternative hypothesis that 2000-2010 is less than 2010-2023: *t* = *−*3.67, *p* = 0.00019).

**Figure 2:**
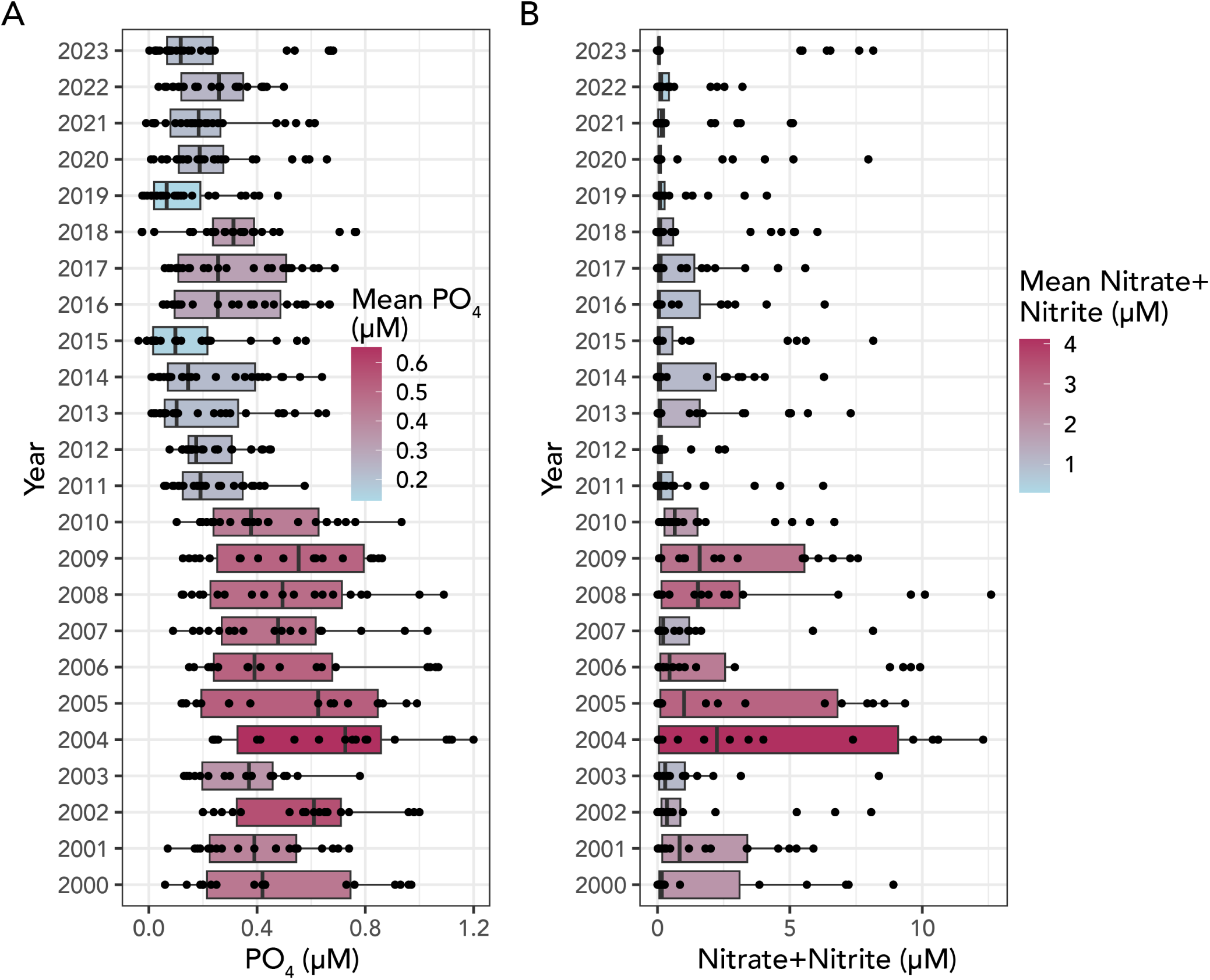
Shift in orthophosphate and combined nitrate and nitrate from 2000 to 2023 Downward shift in availability of nutrients in Cape Cod Bay (CCB) after 2010. A: The shift in mean orthophosphate availability after 2010 was particularly pronounced, with values below 0.1 *µM* as used in the gene abundance comparison becoming much more frequent following 2010. B: Shift in the availability of combined nitrate and nitrite after 2010; after October 2010, combined nitrate and nitrite was measured rather than individual nitrate and nitrite, so values for nitrate and nitrite N concentration were summed and then averaged across surface depths for sampling points prior to 2010 for intercomparability.

**Figure 3:**
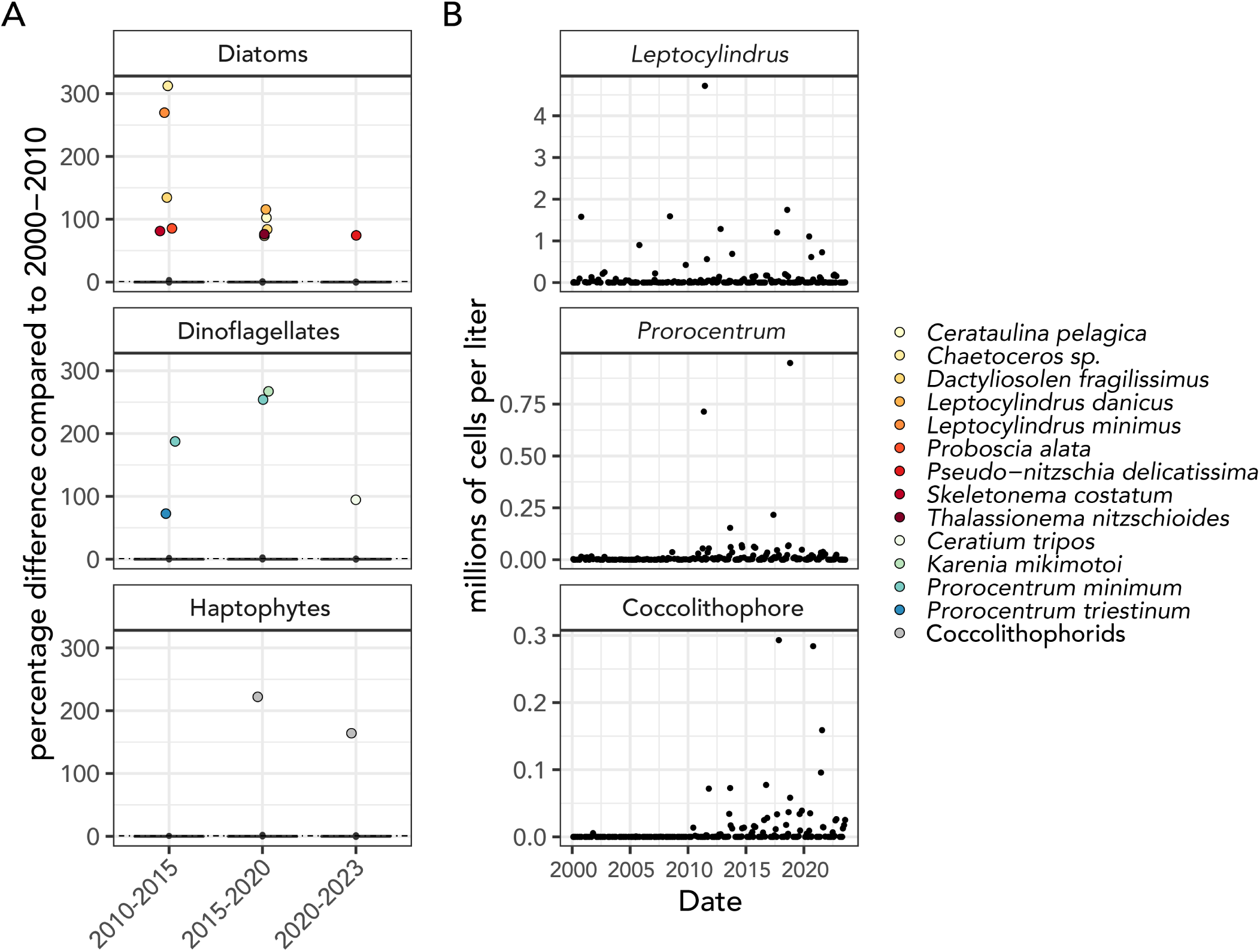
Changes in taxon abundance from counts from pre-2010. Shifts in abundance of taxa within major functional groups in 2010-2015, 2015-2020, and 2020-2023 as compared to 2000-2010 (“baseline”). A: For diatom, dinoflagellate, and haptophyte taxa, groups that had more than a 70% change in abundance in 2010-2015, 2015-2020, or 2020-2023 (*x*-axis) are shown as colored points, while the boxplots show the overall distribution of low percentage differences. B: Time-series abundances of taxa that were more likely to show significant abundance changes in the latter part of the time series; top: *Leptocylindrus*, which only showed overall abundance differences in 2010-2015 and 2015-2020, but showed some seasonal changes in 2020-2023 (Supplementary Figure 9). Middle: *Prorocentrum*, which showed most significant differences in abundance in 2015-2020 alongside *Karenia mikimotoi*. Bottom: general coccolithophore abundance, which became nonzero post-2010.

### 3.2 Nutrient conditions predict phytoplankton trends

Changes in abundances of certain phytoplankton groups appear to be correlated with decreasing trends in micromolar phosphate concentrations (Figure 2). The summer concentration of coccolithophores is increasing in conjunction with an overall decrease in phosphate availability in CCB (Figure 2; Supplementary Figure 9). 6 of 9 (66.6%) sampling points with a recorded concentration of 1 *×* 10^4^ cells L*^−^*^1^ or higher (10 of 13 (76.9%) of those with 5 *×* 10^3^ cells L*^−^*^1^ or greater concentration) had relatively low measured orthophosphate concentration below 0.25 *µ*mol. Coccolithophores both become measurable after the decrease in average orthophosphate after 2010 and continue to be most abundant at the stations and seasonal timepoints at which orthophosphate concentrations are lowest, which tends to be either in midsummer (June or July) or in early fall (September or October; Figure 3B, Supplementary Figure 10).

Within the metatranscriptomic data, the abundance of transcripts annotated as family Noelaerhabdaceae containing *G. huxleyi* had a Spearman rank correlation coefficient of −0.63 (adjusted *p <* 10*^−^*^9^) between environmental orthophosphate concentration and the relative abundance of coccolithophores in the metatranscriptomes as estimated by Salmon (Patro et al., 2017) and reported in transcripts per million (TPM) prior to normalization to group abundance (Figure 4). Metatranscriptomes hence recapitulated the observation that low orthophosphate concentrations tended to correspond to high microscopic counts of coccolithophore abundance, and indicate that high coccolithophore abundance was accompanied by metabolic activity. Though transcript abundance is not expected to correlate directly to cell counts, agreement between metatranscriptomes and counts was generally acceptable for groups of interest (Supplementary Table 1). Coccolithophore abundance was negatively correlated to both combined nitrate and nitrite concentration (correlation= *−*0.38; adjusted *p* = 0.017) and to silica concentration (correlation= *−*0.46; adjusted *p <* 10*^−^*^3^). Coccol-ithophore abundance was correlated positively to temperature (Spearman rank correlation: 0.76; *p <* 10*^−^*^14^). Additional families were selected to compare with family Noelaerhabdaceae and order Prorocentrales on the basis of their abundance patterns. Orders Cryptomonadales and Thalassiosirales tend to have highest abundance in winter and spring, in contrast to coccolithophores and summer dinoflagellates. These groups specifically both had relatively low abundance in July 2021 when Noelaerhabdaceae was most relatively abundant (Figure 11). Because it tends to be most abundant in midsummer, diatom order Leptocylindrales was also used for comparison. Coccolithophores had the strongest negative Spearman correlation with phosphate concentrations, but dinoflagellates (Prorocentrales; correlation = *−*0.51, adjusted *p <* 10*^−^*^4^) and the diatom order Leptocylindrales (correlation= *−*0.49; adjusted *p <* 10*^−^*^3^) were also associated with low phosphate. The abundance of the diatom order Thalassiosirales and cryptophyte order Cryptomonadales were instead positively correlated with orthophosphate (Thalassiosirales: correlation = 0.45, adjusted *p <* 10*^−^*^3^; Cryptomonadales: correlation = 0.64, adjusted *p <* 10*^−^*^8^) and negatively correlated to temperature (Thalassiosirales: correlation = 0.48, adjusted *p <* 10*^−^*^3^; Cryptomonadales: correlation = 0.78, adjusted *p <* 10*^−^*^15^). Additional correlations and their significance values are shown in Figure 4.

**Figure 4:**
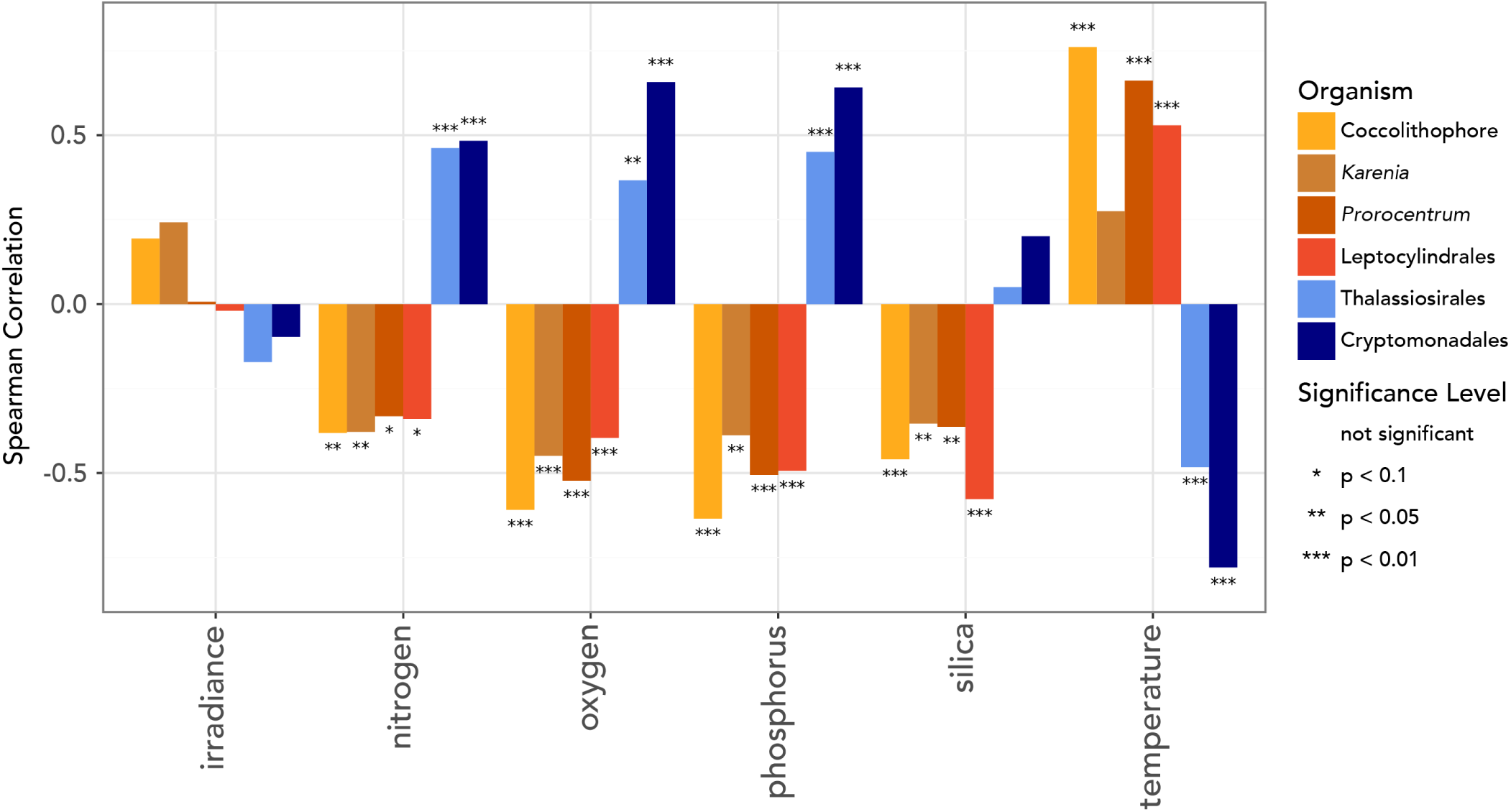
Spearman rank correlation between abundance and environmental variables for important groups. Spearman *ρ* rank correlation coefficient values for comparisons with *p*-value less than 0.05 between abundances of different taxonomic families and orthophosphate concentrations, after Bonferroni adjustment for multiple comparisons. In order to be shown, a taxonomic group also had to have *≥* 10^4^ summed total TPM abundance across all samples.

**Figure 5:**
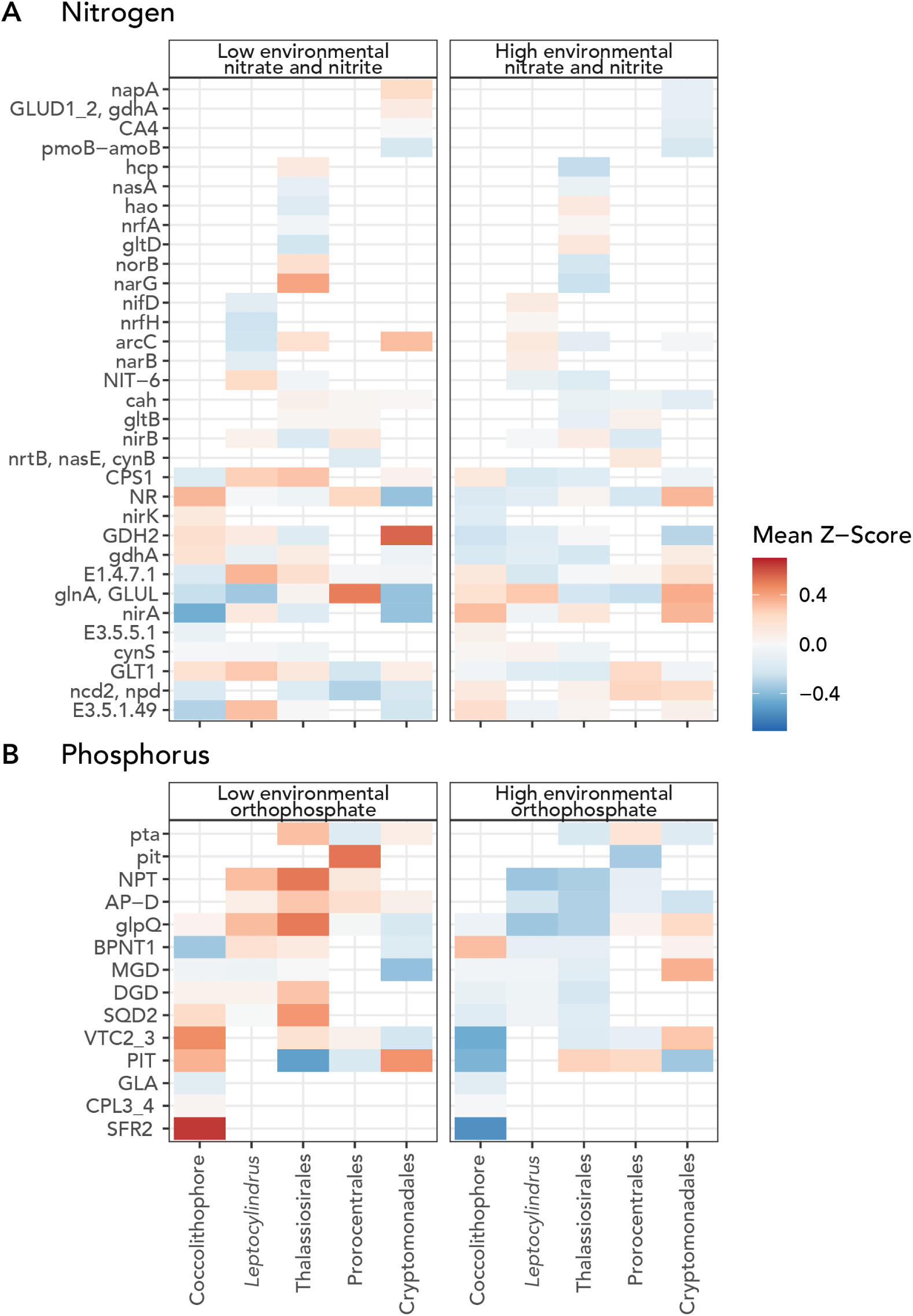
Correlation of N and P-associated genes. For each of 5 taxonomic groups, abundance of gene transcripts associated with nitrogen and phosphorus relative to the environmental availability of the total of nitrate and nitrate and orthophosphate, respectively. A: Nitrogen use-associated genes and their average *Z*-scores under high vs. low summed nitrate and nitrite concentrations. “Low” nitrate and nitrite was defined as 0.1 *µM* . B: Phosphorus use-associated genes and their average *Z*-scores under high vs. low orthophosphate concentrations. “Low” orthophosphate was defined as 0.05 *µM* .

### 3.3 Phosphate-related genes differentiate phytoplankton metabolic strategies

While phosphate-related genes (Supplementary Table 2) tended to be correlated with low environmental phosphate across taxa (number samples = 27), many of the specific genes used to respond to phosphate limitation were exclusively used by particular taxa.

#### 3.3.1 Identifying genes associated with very low phosphate availability

In coccolithophores, three genes related to phosphate limitation showed higher expression normalized to total coccolithophore abundance across the time series under low (*<* 0.1 *µ*M) orthophosphate: *VTC4* (KEGG: K27692), a vacuolar transporter chaperone (VTC) complex gene involved in algal polyphosphate storage (Cliff et al., 2023) (*t* = 4.84, adjusted *p <* 10*^−^*^4^), and *PIT* (KEGG: K14640), a sodium-dependent phosphate transporter (*t* = 4.32, adjusted *p <* 10*^−^*^3^), and a phospholipid:diacylglycerol acyltransferase (KEGG: K00679) shown previously to be involved in microalgal nutrient limitation response (Sah et al., 2024) (*t* = 8.66, adjusted *p <* 10*^−^*^8^). One additional significant orthophosphate-responsive gene was *SFR2* (KEGG: K21362), a galactolipid galactosyltransferase implicated in response to freezing in plants (Thorlby, Fourrier, and Warren, 2004), but part of the beta-glucosidase family; other members of which have been shown to be phosphate starvation-inducible (Malboobi and Lefebvre, 1997), and further may be involved in a physiological shift from phospholipids to galactolipids under P-limitation (Kelly, Froehlich, and Dörmann, 2003) (*t* = 3.92, *p* = 0.001). The dinoflagellate order Prorocentrales had no significant pairwise differences between phosphate-related gene abundance at low (*<* 0.1 *µ*M) vs. high orthophosphate concentrations. In Leptocylindrales, three phosphate-related genes were significantly associated with low environmental orthophosphate: *glpQ* (*t* = 4.27, adjusted *p* = 0.00033), a phosphate transport system protein *pstS* (KEGG: K02040; *t* = 3.86, adjusted *p* = 0.00088), and *NPT*, a sodium-dependent phosphate co-transporter (KEGG: 14683; *t* = 3.50, adjusted *p* = 0.0039). The sodium-dependent phosphate transporter (*PIT*; KEGG: K014640) was more highly expressed under low environmental orthophosphate in cryptophytes (*t* = 4.21, adjusted *p* = 0.000038). In the diatom Thalassiosirales, a sodium-dependent phosphate cotransporter (KEGG: K14683; *t* = 3.24, adjusted *p* = 0.00059), an AP-3 complex subunit involved in inositol phosphate regulation (Hao et al., 1997) (KEGG: K12398; *t* = 3.04, adjusted *p* = 0.008), and *glpQ* (KEGG: K01126; *t* = 3.41, adjusted *p* = 0.00039) were all more highly expressed when orthophosphate was low.

#### 3.3.2 Coccolithophores have a comparatively high number of genes correlated with orthophosphate concentration

We also calculated correlations between normalized expression (*Z*-scores) and orthophosphate concentration, and found that coccolithophores had the highest number of genes significantly negatively correlated to orthophosphate (*n* = 2058 of 5635), compared to *n* = 1903 of 5427 for Leptocylindrales, *n* = 1342 of 3999 for Prorocentrales, *n* = 1185 of 7321 for Thalassiosirales, and *n* = 628 of 6134 for Cryptomonadales (Figure 7B; Leptocylindrales, Prorocentrales, and coccolithophores all have a significantly higher number of correlated genes). In coccolithophores, *VTC4* (*ρ* = *−*0.55, adjusted *p* = 2.2e*^−^*^6^), *PIT* (*ρ* = *−*0.53, adjusted *p <* 10*^−^*^5^), the phospholipid:diacylglycerol acyltransferase (*ρ* = *−*0.78, adjusted *p <* 10*^−^*^13^), and *SFR2* (*ρ* = *−*0.61, adjusted *p <* 10*^−^*^7^) all had significant correlations to low environmental orthophosphate, alongside the sulfoquinovosyltransferase (*SQD2*; KEGG: K06119) enzyme needed to convert a precursor molecule to a sulfolipid (*ρ* = *−*0.27, adjusted *p* = 0.03), *DGD* (KEGG: K09480), a digalactosyldiacylglycerol synthase also implicated in galactolipid synthesis (Härtel, Dörmann, and Benning, 2000; Kelly, Froehlich, and Dörmann, 2003; Kelly and Dörmann, 2002) (*ρ* = *−*0.36; *p* = 0.004), *glpQ* (KEGG: K01126), a glycerophosphoryl diester phosphodiesterase involved in phospholipid metabolism (*CPL3_4* (KEGG: K18999); *ρ* = *−*0.42, adjusted *p <* 10*^−^*^3^, and a phosphatase that phosphorylates the C-terminal domain of RNA polymerase II (*ρ* = *−*0.47, adjusted *p* = 1.0e*^−^*^4^). Two Prorocentrales genes had significant time series correlations with environmental orthophosphate: a sodium-dependent phosphate cotransporter (KEGG: K14683; *ρ* = *−*0.29, adjusted *p* = 0.021) and diacylglycerol galactosyltransferase (*MGD*; KEGG: K03715) shown to be involved in lipid remodeling in response to phosphate starvation in plants (*ρ* = *−*0.33, adjusted *p* = 0.0083). All three of the Leptocylindrales phosphate-related genes that were more highly expressed under low phosphate were also correlated to phosphate concentrations (*glpQ*: *ρ* = *−*0.47, adjusted *p* = 0.0001; *pstS*: *ρ* = *−*0.40, adjusted *p* = 0.0001; *NPT*: *ρ* = *−*0.45, adjusted *p* = 0.00018), in addition to *BPNT1* (KEGG: K01082), an enzyme which converts 3’-phosphoadenosine-5’-phosphate (PAP) to adenosine-5’-phosphate (AMP) and inorganic phosphate (Pi) (*ρ* = *−*0.33, adjusted *p* = 0.0075) and alkaline phosphatase D (KEGG: K01113; *ρ* = *−*0.33, adjusted *p* = 0.00089). Like Leptocylindrales, all three of the Thalassiosirales genes with higher expression when phosphate was low were also correlated to low environmental orthophosphate across the timeseries (sodium-dependent phosphate cotransporter: *ρ* = *−*0.70, adjusted *p* = 1.77e*^−^*^10^; AP-3 subunit: *ρ* = *−*0.36, adjusted *p* = 0.0032; *glpQ*: *ρ* = *−*0.55, adjusted *p* = 2.0e*^−^*^6^), in addition to VTC4 (KEGG: K27692; *ρ* = *−*0.29, adjusted *p* = 0.02), *DGD* (KEGG: K09480; *ρ* = *−*0.39, adjusted *p* = 0.0017), and alkaline phosphatase D (KEGG: K01113, *ρ* = *−*0.48, adjusted *p <* 10*^−^*^4^).

#### 3.3.3 Differential expression analysis between July 2021 and 2023 highlights between-summer differences

For all taxonomic groups, we conducted differential expression analysis between the measurements at all three stations (F01, F02, and F29) in July of 2021 as compared to July of 2023 (Figure 6A). In July of 2021, Prorocentrales, coccolithophores, and were relatively abundant, but coccolithophores were especially relatively abundant (Supplementary Figure 10). Genes that were upregulated in July 2021 compared to July 2023 at all stations for coccolithophores included a sodium-dependent phosphate transporter (*PIT*; KEGG: K14640; log-fold change 4.15; adjusted *p <* 10*^−^*^29^), *SFR2* (KEGG: K21362; log-fold change 3.19; adjusted *p <* 10*^−^*^17^), and UDP-sulfoquinovose synthase (KEGG: K06118; log-fold change 1.68; adjusted *p* = 0.013) and sulfoquinovosyltransferase (KEGG: K06119; log-fold change 2.23; adjusted *p <* 10*^−^*^3^), the sequential (*SQD1* and *SQD2*, respectively) enzymes needed to convert a precursor molecule to a sulfolipid, *glpQ* (KEGG: K01126; log-fold change 2.31; adjusted *p* = 0.017), alkaline phosphatase (KEGG: K01077; log-fold change 1.17; adjusted *p* = 0.014), phospholipid:diacylglycerol acyltransferase (KEGG: K00679; log-fold change 2.81, adjusted *p* = 0.0088), vacuolar transport protein VTC2_3 (KEGG: K27692; log-fold change 2.46, adjusted *p <* 10*^−^*^8^), and a phosphate transporter *pstS* (KEGG: K02040; log-fold change 3.90; adjusted *p <* 10*^−^*^6^). Prorocentrales genes upregulated in July 2021 compared to July 2023 included a sodium-dependent phosphate cotransporter (KEGG: K14683; log-fold change 4.03, adjusted *p <* 10*^−^*^4^), VTC2_3 (KEGG: K27692; log-fold change 4.77, adjused *p <* 10*^−^*^4^), and *DGD* (KEGG: K09480; log-fold change: 4.26, adjusted *p* = 0.0029), a digalactosyldiacylglycerol synthase also implicated in galactolipid synthesis (Härtel, Dörmann, and Benning, 2000; Kelly, Froehlich, and Dörmann, 2003; Kelly and Dörmann, 2002). In Leptocylindrales, only one functional gene–a sodium-dependent phosphate cotransporter (KEGG: K14683; log-fold change 4.58, adjusted *p <* 10*^−^*^29^)–was upregulated in July 2021 compared to July 2023. Cryptomonadales had two phosphate-related genes upregulated in July 2021 compared to July 2023: alkaline phosphatase D (KEGG: K01113; log-fold change 4.84; adjusted *p* = 0.008), and UDP-sulfoquinovose synthase (*SQD1*; KEGG: K06118; log-fold change 4.12; adjusted *p* = 0.02). An inorganic phosphate transporter (KEGG: K03306; log-fold change −3.81; adjusted *p* = 0.005) was upregulated in July of 2023 compared to July of 2021. Cryptophyte upregulation of this inorganic phosphate transporter in July 2023 might be associated with the slightly elevated orthophosphate concentration in July 2023 as compared to 2021. In order Thalassiosirales, one gene was upregulated in July 2021 compared to July 2023: a sodium-dependent phosphate cotransporter (KEGG: K14683; log-fold change 4.46; adjusted *p <* 10*^−^*^9^). Two phosphate-related genes, a vacuolar transport protein (*VTC2_3*, KEGG: K27692; log-fold change −2.88, adjusted *p* = 0.01) and *glpQ*, glycerophosphoryl diester phosphodiesterase (KEGG: K01126, log-fold change −3.20, adjusted *p* = 0.0022), were upregulated in July 2023 compared to July 2021. All differential expression results for July 2021 as compared to July 2023 are summarized in Figure 6A.

**Figure 6:**
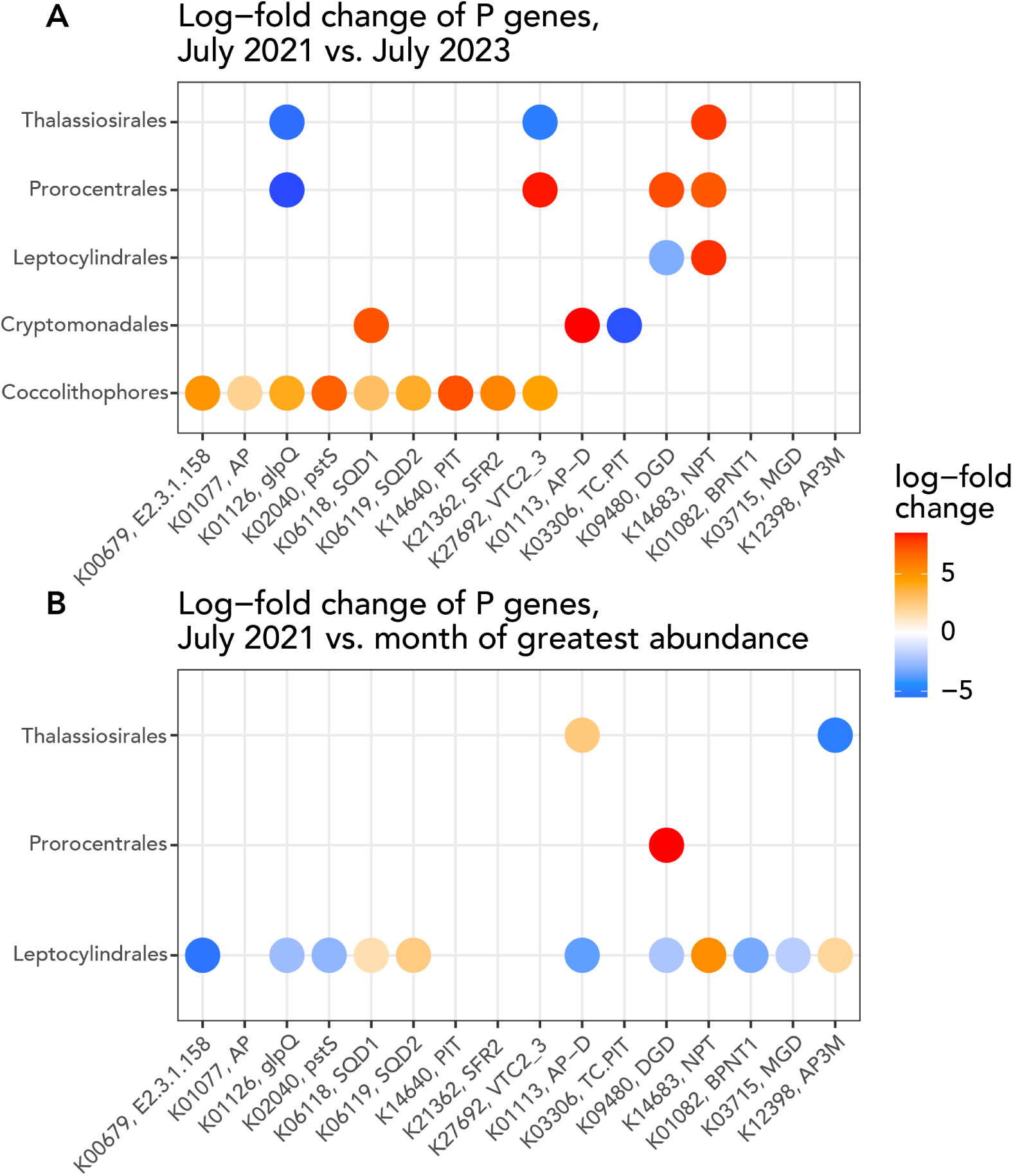
Summary of differential expression results for genes associated with phosphate use. A: significantly differentially-expressed genes with regulation differences assessed in July 2021 as compared to July 2023. B: significantly differentially-expressed phosphate-related genes for groups other than coccolithophores (cryptophytes also tested, but no significant differences in expression were detected) in July 2021 as compared to the month of that organism’s greatest abundance in the three-year time series.

**Figure 7:**
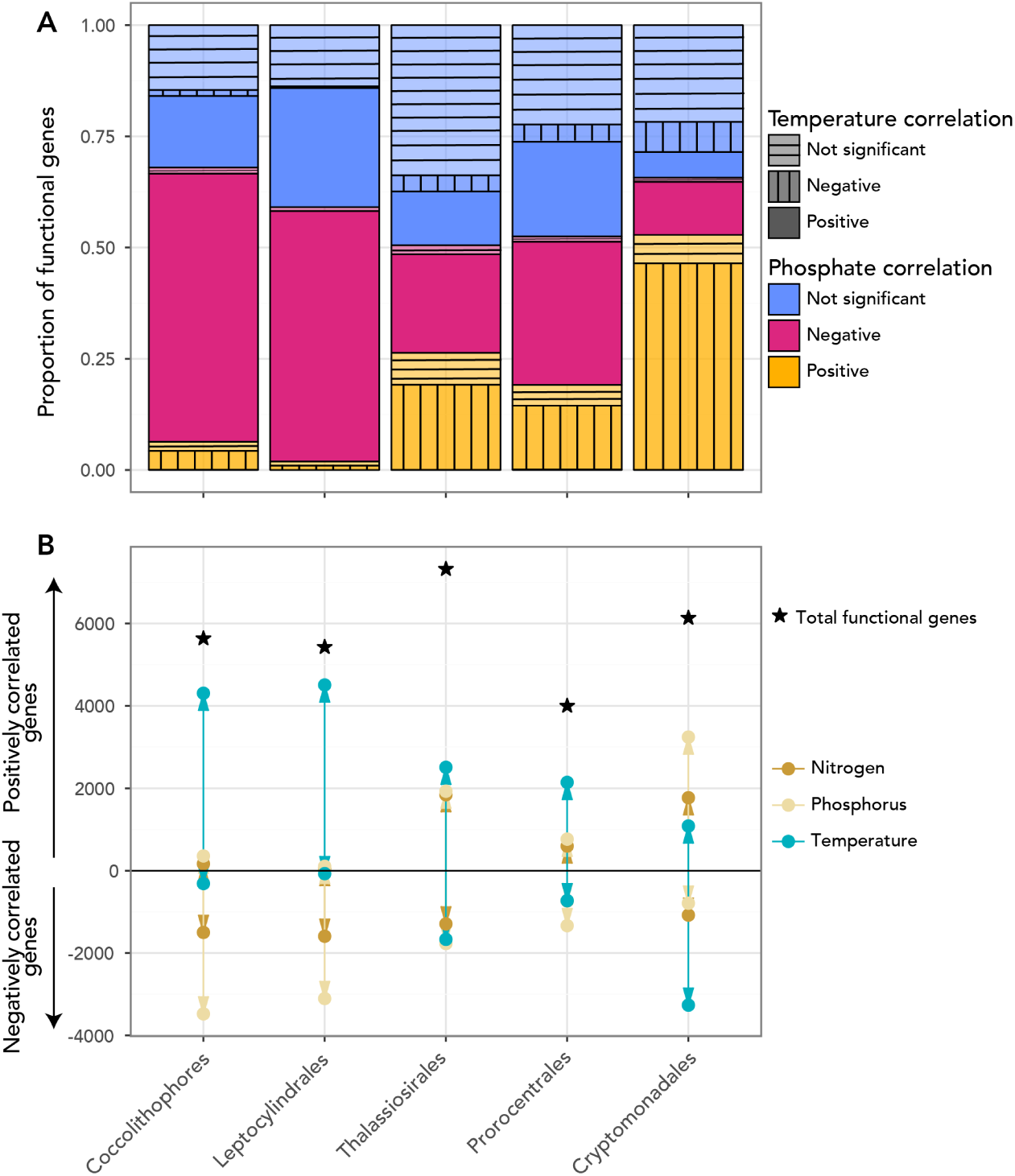
Summary of Spearman correlation results between functionally-annotated genes and environmental conditions (orthophosphate concentration, nitrate/nitrite concentration, and temperature). A: The proportion of identified functional genes within each taxonomic group that was correlated with phosphate (color) compared to temperature (alpha/hatching pattern). The highest number of genes were positively correlated to temperature and negatively correlated to phosphate for coccolithophores, Leptocylindrales, and Prorocentrales, whereas more genes were positively correlated to phosphate and negatively correlated to temperature in cryptophytes and Thalassiosirales. B: Number of functionally-annotated genes positively vs. negatively correlated with nitrate/nitrite, orthophosphate, and temperature (colors); number negatively correlated is shown as a negative count (below the horizontal line at zero), while number positively correlated is shown as a positive count (above the horizontal line at zero). Stars above the positive count points and arrows show the total number of identified functional genes for each group, with the balance of genes not showing significant correlations.

#### 3.3.4 Comparing July 2021 phosphate limitation to the month of highest abundance

For all groups except coccolithophores, we conducted differential expression analysis between July 2021 at F29 and the timepoint at F29 at which each other taxonomic group had highest relative gene expression (May of 2022 for Prorocentrales, July of 2022 for Leptocylindrales, February 2023 for Cryptomonadales, and March 2021 for Thalassiosirales; Figure 6B; Supplementary Figure 12). We compared station F29 between July of 2021 and May of 2022 for Prorocentrales and found one P-related gene significantly upregulated in July of 2021: *DGD* (KEGG: K09480), a digalactosyldiacylglycerol synthase (log-fold change 8.0, adjusted *p* = 0.03). Comparing station F29 between July 2021 and February 2023 for order Cryptomonadales, we found no phosphate-related genes that were significantly differentially expressed in February 2023 compared to July 2021. Leptocylindrales had four phosphate-related genes upregulated in July 2021 compared to July 2022: UDP-sulfoquinovose synthase (*SQD1*, KEGG: K06118; log-fold change 1.39, adjusted *p* = 0.006) and *SQD2* (KEGG: K06119; log-fold change 4.42, adjusted *p <* 10*^−^*^4^), an AP-3 complex gene (K12398; log-fold change 1.79, adjusted *p* = 0.0077), and a sodium-dependent phosphate cotransporter (KEGG: K14683; log-fold change 4.94, adjusted *p <* 10*^−^*^17^). Conversely, seven phosphate-related genes were upregulated in July of 2022 compared to July of 2021: *BPNT1* (KEGG: K01082), an enzyme which converts 3’-phosphoadenosine-5’-phosphate (PAP) to adenosine-5’-phosphate (AMP) and inorganic phosphate (Pi; log-fold change −3.0, adjusted *p <* 10*^−^*^13^), a phospholipid:diacylglycerol acyltransferase (KEGG: K00679; log-fold change −5.0, adjusted *p* = 0.0026), alkaline phosphatase D (KEGG: K01113; log-fold change −3.41, adjusted *p <* 10*^−^*^11^), a phosphate transport system protein *pstS* (KEGG: K02040; log-fold change −2.59, adjusted *p <* 10*^−^*^6^), *glpQ* (KEGG: K01126), a glycerophosphoryl diester phosphodiesterase involved in phospholipid metabolism (log-fold change −2.32, adjusted *p* = 0.027), *DGD* (KEGG: K09480; log-fold change 2.0, adjusted *p <* 10*^−^*^6^), and diacylglycerol galactosyltransferase (*MGD*; KEGG: K03715; log-fold change −1.72, adjusted *p <* 10*^−^*^11^). In Thalassiosirales, one phosphate-related gene was significantly upregulated in March 2021 compared to July 2021: the AP-3 complex gene (K12398; log-fold change −4.70, adjusted *p* = 0.013), while alkaline phosphatase D (KEGG: K01113; log-fold change 2.34, adjusted *p* = 0.022) was upregulated in July 2021 compared to March 2021. All differential expression results are summarized in Figure 8 , highlighting the investment of coccolithophores in a suite of phosphate limitation-related genes in July of 2021 and demonstrating that use of these genes is not likely correlated to cell abundance or growth rates for other taxa. All differential expression results are summarized in Figure 6B.

**Figure 8:**
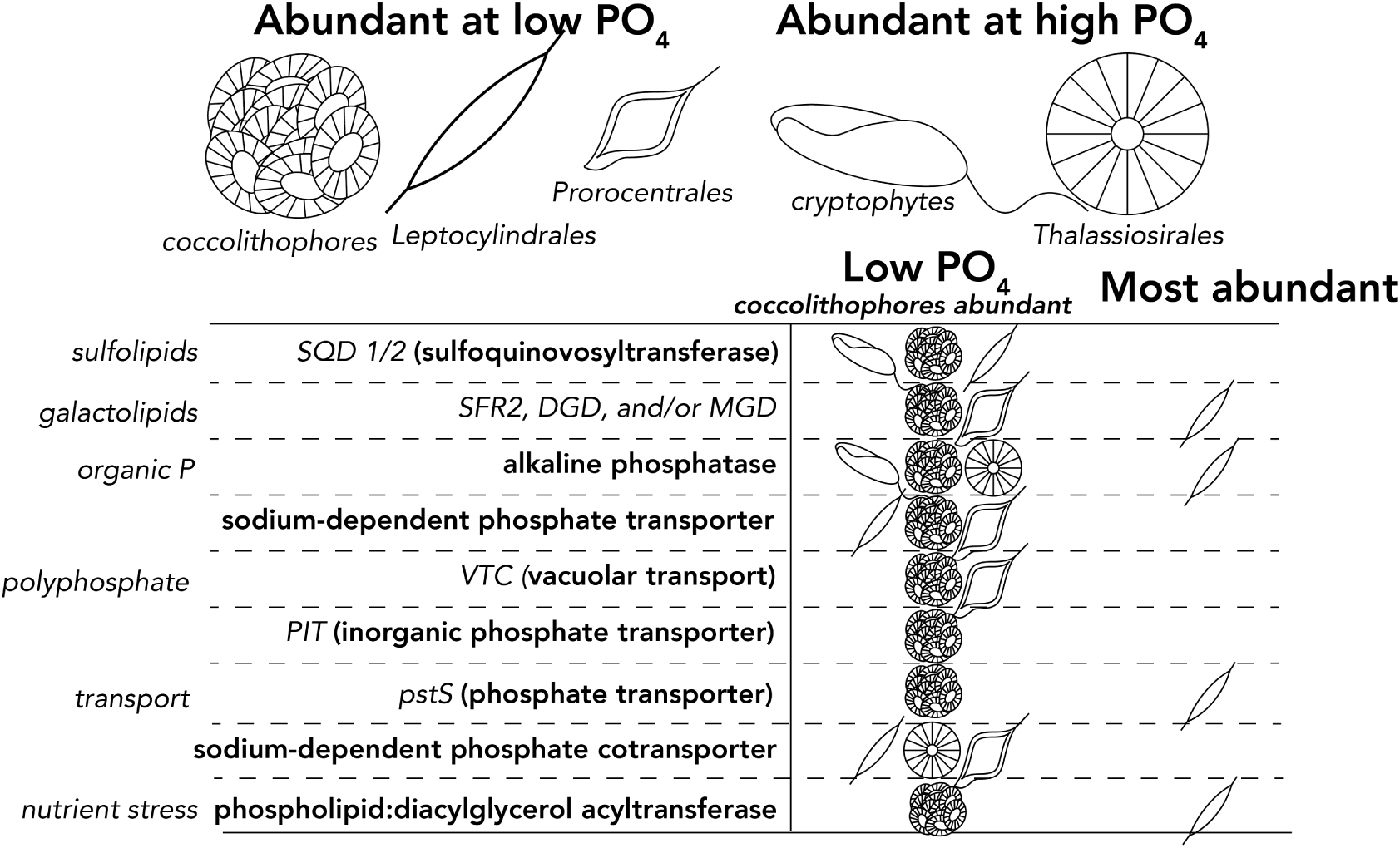
Synthesis of observations of phytoplankton investment in phosphate-related genes according to environmental conditions. The low phosphate (PO_4_) column corresponds to July of 2021, when coccolithophores were most abundant. Genes were counted as more highly expressed in July of 2021 when comparisons were made either to July of 2023 or to the month in which the organism of interest was most abundant, for all except coccolithophores (March of 2021 for Thalassiosirales, May of 2022 for Prorocentrales, July of 2022 for Leptocylindrales, and February of 2023 for Cryptomonadales/cryptophytes).

#### 3.3.5 Genes with correlations to multiple environmental variables

We further clustered genes within each of the five taxa by the correlations of individual gene abundances with nitrate, phosphate, and temperature. We identified genes contained in clusters that were, e.g., similarly correlated to phosphate in multiple taxa. We were further interested in genes that were correlated with one environmental variable and not others, e.g., with temperature but not with some nutrient, so we conducted a second clustering of the matrix of correlations with environmental variables for each functional gene. We discovered that a lysophospholipase gene (KEGG: K06129) was positively correlated to phosphate in Prorocentrales while negatively to temperature, but negatively correlated to phosphate in coccolithophores. This enzyme is involved in freeing up phosphate from phospholipids as sulfolipids and galactolipids are being synthesized, but also in the Lands cycle in creating new phospholipid structures to stabilize membranes under e.g., temperature limitation. In coccolithophores, this gene is one of the few genes not significantly correlated with temperature, but significantly negatively correlated with phosphate, meaning that its investment increases when phosphate becomes limiting and not exclusively under thermal limitation. Clustering all functional genes according to their correlation with these three environmental conditions highlights commonalities between summer-associated taxa (coccolithophores, Leptocylindrales, and Prorocentrales) as compared to winter and spring-associated taxa.

## 4 Discussion

To investigate mechanisms underlying the success of emerging taxa in coastal ecosystems under climate change, we incorporated metatranscriptomic sequencing into a long-term oceanographic survey program to attribute underlying transcriptional explanations for observed increase in coccolithophores and species of the genus *Prorocentrum* in an important coastal ecosystem. Cape Cod Bay (MA, USA; CCB) is an important recreation and fishing area that constitutes no less than $1.5 billion of the local region’s total economic productivity (Costa and Hughes, 2012), and impacts the Boston metropolitan area. Hence, water quality research is critical in CCB, and we can consider CCB a prototypical ecosystem for future shifts in coastal phytoplankton dynamics. Our *Z*-score method applied to time series metatranscriptomes from this vulnerable coastal region considers metabolism across the full range of conditions measured in the time series rather than comparing only one individual sampling point to another at a time. Our findings are further supported by differential expression analysis, which permits statistical evaluation of pairwise comparisons between conditions. We found similarities between organisms that were most abundant in summer, nutrient-depleted conditions (including coccolithophores, dinoflagellates, and groups of diatoms like Leptocylindrales), but distinctive features that differentiated functional groups. The emergence of mixotrophic, generalist dinoflagellates like *Prorocentrum* (Telesh, Schubert, and Skarlato, 2016) in alternate months and years in the dataset emphasizes the development of diverse new niches in a changing coastal ecosystem. These diverse niches shift ecosystem trophic state and expected nutrient export via transition between mixotrophs and phototrophs with unique ecological traits, such as toxin production or inorganic carbon use. Overall, our results illuminate long-term changes in relevant coastal phytoplankton taxa and potential drivers of these changes.

Changes in abundance of the formerly rare taxa that we targeted in this study have far-reaching implications for biogeochemistry and human health. *G. huxleyi* and other coccolithophores are known for high affinity for orthophosphate (Paasche, 2001; Paasche, 1998), but are also expected to thrive in oligotrophic open ocean conditions (Paasche, 2001). Though *G. huxleyi* ordinarily has a negligible coastal impact, relatively large coastal blooms have been recently observed in concert with higher temperatures, in response to a diatom bloom (Matson et al., 2019) and even as a consequence of a competitive advantage offered by oil exposure (Ladd et al., 2018). *G. huxleyi*’s metabolic traits may allow it to thrive in an ephemeral coastal niche characterized by nutrient depletion and enabled by the intensive nutrient requirements and large export (Smith Jr et al., 2021) of colonial blooms like those formed by *Phaeocystis pouchetii* or fast-sinking blooms formed by diatoms. Warming water temperatures may accelerate the timeline of the spring bloom and prolong the period of nutrient limitation. Phosphate starvation may also increase calcification (Paasche, 1998) and change membrane characteristics (Shemi et al., 2016). *G. huxleyi*’s mixotrophic potential (Godrijan, Drapeau, and Balch, 2020) may also enable occasional growth of *G. huxleyi* in highly nutrient-depleted environments, deeper waters with low-light conditions, seasonally lower light availability, or shading (self- or due to high-density blooms of other organisms) (Godrijan, Drapeau, and Balch, 2020; Linge Johnsen and Bollmann, 2020). We observed increased coccolithophore inorganic phosphate transport gene expression and modestly increased expression of alkaline phosphatase genes (Martin et al., 2014) in response to field phosphate limitation. We also found evidence for a significant role of phospholipid substitution with galactolipids and sulfolipids among coccolithophores in CCB, which provides a mechanism by which phytoplankton can take advantage of ephemeral phosphate limitation in coastal environments. The observation that *G. huxleyi* may use lipid substitution to tolerate profoundly low available orthophosphate is supported by existing literature documenting that *G. huxleyi* is likely to perform lipid substitution to cope with limited phosphate in the laboratory (Van Mooy et al., 2009; Shemi et al., 2016).

The genus *Prorocentrum* is a group of dinoflagellates known to be a constituent of summer and fall phytoplankton blooms in CCB (Hunt et al., 2010). Dinoflagellates like *Prorocentrum* may have human health impacts through the production of toxins (Telesh, Schubert, and Skarlato, 2016), may become more abundant under climate change conditions (Boivin-Rioux et al., 2021), and increased temperature may increase cellular chlorophyll a content (Fu et al., 2008). *Prorocentrum* has also been documented to be invasive in other systems such as the Baltic Sea (Telesh, Schubert, and Skarlato, 2016). Nutrient limitation may induce mixotrophy in *Prorocentrum*, resulting in capacity to compete under nutrient-limited conditions (Johnson, 2015). Other toxin-producing dinoflagellates have also been observed recently in the Gulf of Maine region, including characteristic *Alexandrium* blooms (Anderson, 1997) and new blooms of *Karenia*. The toxic dinoflagellate *Karenia mikimotoi* was first documented in CCB in 2017 (Scully et al., 2022). *K. mikimotoi* is responsible for large, economically-damaging blooms that can trigger hypoxic conditions in deeper waters when they sink (Li et al., 2019; Scully et al., 2022). *K. mikimotoi* concentrations have been linked to low salinity and high silicate and nutrient concentrations, and *K. mikimotoi* growth frequently excludes diatoms (Barnes et al., 2015). Low light levels have also been linked to the success of *K. mikimotoi* given stable environmental conditions and sufficient nutrient input (Su et al., 2024; Guangmao and Shufeng, 2018). Though *K. mikimotoi* blooms were documented from 2017-2019, cell numbers were relatively low in CCB from 2021-2023 when we conducted our metatranscriptomic sampling. Combined with challenges of *Karenia* and other dinoflagellate transcriptomics due to the prevalence of post-transcriptional modifications (Morey et al., 2011), we decided to focus only on *Prorocentrum* in this study. However, the observations of recent blooms of *K. mikimotoi* highlight shifts in major constituents of the phytoplankton community in CCB and the emergence of dinoflagellate taxa that threaten human health. The molecular data highlight that Prorocentrales also leverages the galactolipid substitution mechanism to persist under unseasonably low phosphate concentrations in midsummer, contributing to its persistence in CCB.

We used diatoms in the order Thalassiosirales and cryptophytes as winter comparison taxa to coccolithophores and dinoflagellates correlated with summer warm water temperatures and nutrient limitation. General changes in galactolipid and sulfolipid metabolism were shared between coccolithophores and diatoms in the order Leptocylindrales under phosphate limitation (including in July of 2022 when Leptocylindrales was relatively abundant and phosphate was seasonally scarce). However, we did not observe signatures of the galactolipid mechanism in Thalassiosirales diatoms or in cryptophytes. The absence of this mechanism in cryptophytes may be in part explained by the ability of some cryptophyte taxa to acquire phosphorus via mixotrophy (Stoecker et al., 2017).

In summary, we leveraged a monthly metatranscriptomic time series coupled to long-term count and environmental physical and chemical sampling to identify the molecular mechanisms responsible for the success of diverse phytoplankton taxa in a coastal ecosystem. Our results support the role of lipid substitution in allowing phytoplankton to persist under phosphate limitation and highlight the strategies for phosphate import and use that enable some unusual phytoplankton taxa to gain a competitive advantage. These results help contextualize shifts in coastal phytoplankton abundance as anthropogenic change results in simultaneous localized change in temperature and inorganic nutrient availability.

## Author contributions

MMB initiated the idea for the metatranscriptomic time-series and designed the protocol for sampling, with input from HA. AC led field sampling and led processing of water quality samples. MMB and AIK led the metatranscriptomic sampling protocol with assistance from SKS, AC, and HA. AIK wrote the paper with collaboration from MMB. HA secured funding and supervised the research; AIK secured additional funding with supervision by HA and MJF. MAF led satellite data analysis and provided direction for data interpretation. SKS and MMB led molecular sample processing. All co-authors contributed to review and editing of the draft.

## Acknowledgments

We gratefully acknowledge McCaela Acord and Alese Schofield for assistance with field sampling and sample processing, and Marc Costa, captain of the R/V Shearwater and of marine operations at the Provincetown Center for Coastal Studies. AIK was supported by a Postdoctoral Fellowship and MAF was supported by a Faculty Fellowship from the Center for Chemical Currencies of a Microbial Planet (C-CoMP; National Science Foundation award OCE-2019589). This material is based upon work supported by the U.S. Department of Energy, Office of Science, Office of Advanced Scientific Computing Research, Department of Energy Computational Science Graduate Fellowship under Award Number DE-SC0020347, under which AIK was supported. MMB was supported by a Simons Foundation Postdoctoral Fellowship in Marine Microbial Ecology (award #874439). HA was supported by a Simons Foundation Early Career Investigator in Aquatic Microbial Ecology and Evolution Award (award #931886). The Simons Collaboration on Computational Biogeochemical Modeling of Marine Ecosystems (CBIOMES) supported MJF and AIK (award #549931). A Grassle Fund grant from Academic Programs at Woods Hole Oceanographic Institution provided funding for this research.

## Supporting information

Additional supporting information may be found in the online version of the article at the publisher’s website.

### A Supplementary Text

#### A.1 Spearman correlation between metatranscriptomic TPM and microscopic counts

Spearman correlations were calculated using the cor.test function in R and were constructed by aggregating counts at each taxonomic level and comparing to TPM relative abundance estimates. Results are summarized in Table 1.

**Table 1:**
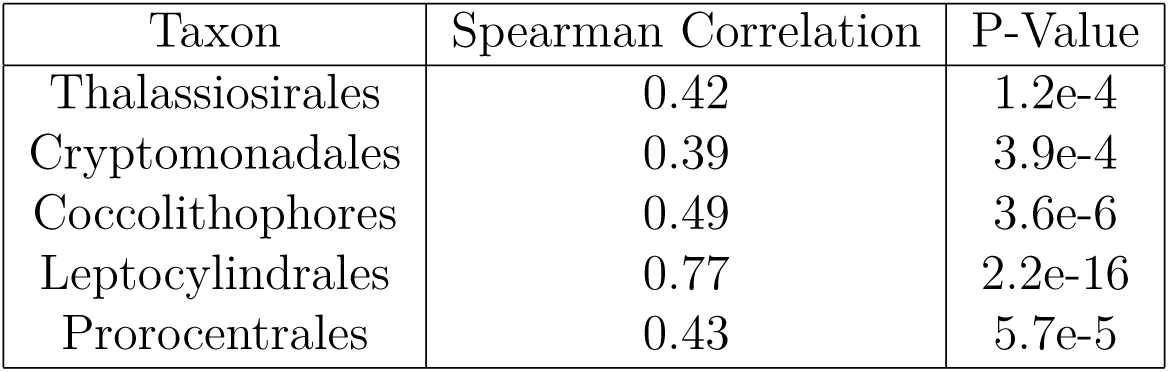
Spearman correlations between summed abundance of each group divided by total counts against TPM relative abundance estimate from metatranscriptomes.

#### A.2 Other organisms affected by changing nutrient patters in Cape Cod Bay

Recurrent and potentially more prevalent seasonal blooms of *Phaeocystis pouchetii* (Jiang et al., 2014; Borkman et al., 2016; Smith Jr et al., 2021) appear to be responsible for higher rates of uptake of both nitrogen and phosphorus in Massachusetts Bay (Smith Jr et al., 2021), and in CCB have been shown to be associated with high phosphate uptake (Borkman et al., 2016). Despite the ability of *Phaeocystis* to store phosphate (Veldhuis, Colijn, and Admiraal, 1991), *Phaeocystis* blooms were preceded by elevated phosphate concentrations and high nitrate to phosphate ratios in Massachusetts Bay in Borkman et al.’s 2016 study (Borkman et al., 2016). The induction of phosphate depletion by *Phaeocystis pouchetii* blooms in Cape Cod Bay could spur an increase in colony formation close to bloom termination, which would further deplete nutrients and enhance carbon export (Veldhuis and Admiraal, 1987; Smith Jr et al., 2021).

#### A.3 Supplementary Figures

**Figure 9:**
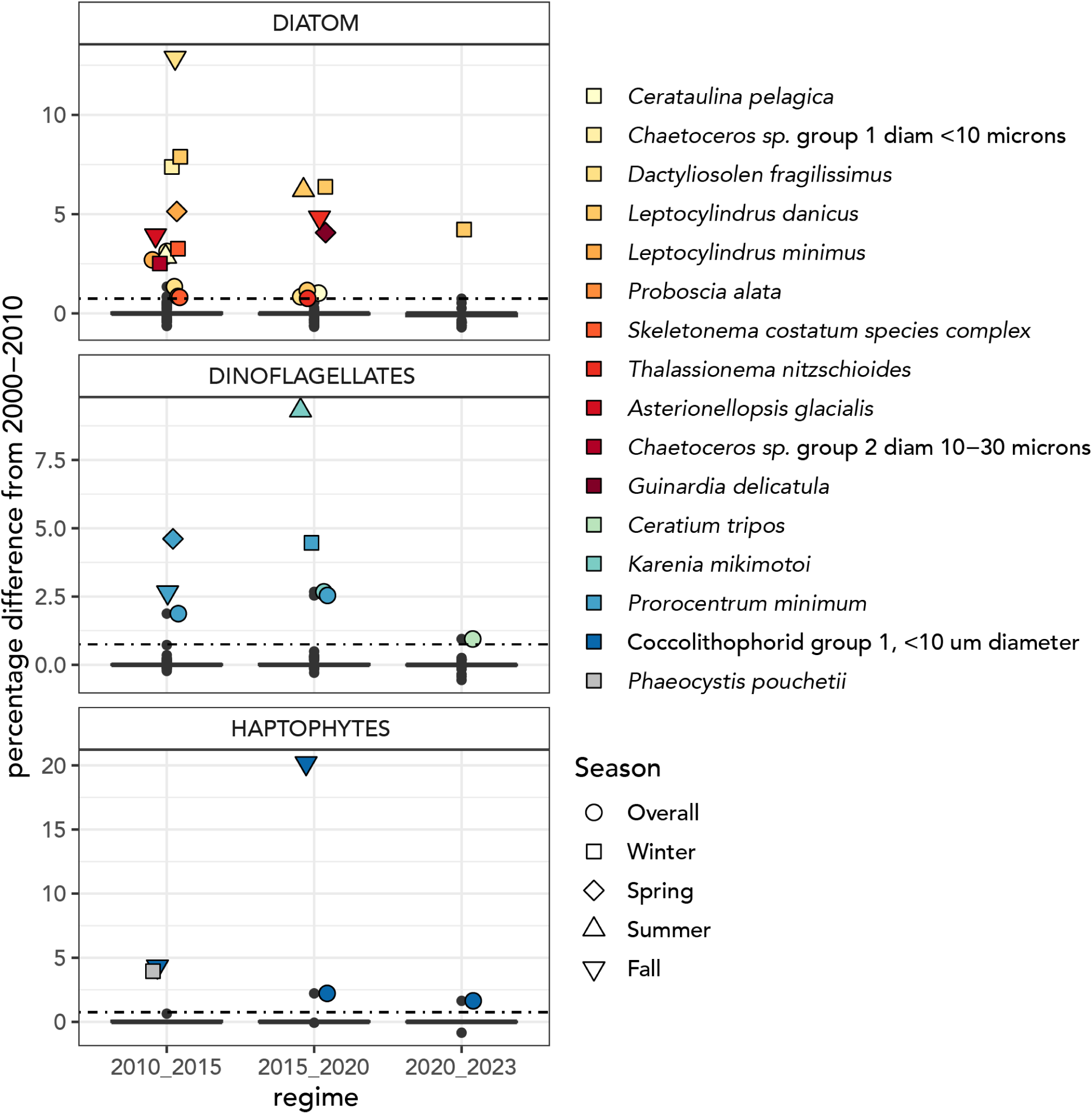
Taxa with significant percentage differences in abundance as compared to 2000-2010 in 2010-2015, 2015-2020, and 2020-2023. Significant changes are shown both overall and by season based on the shape of the marker.

### B Supplementary Tables

**Table 2:**
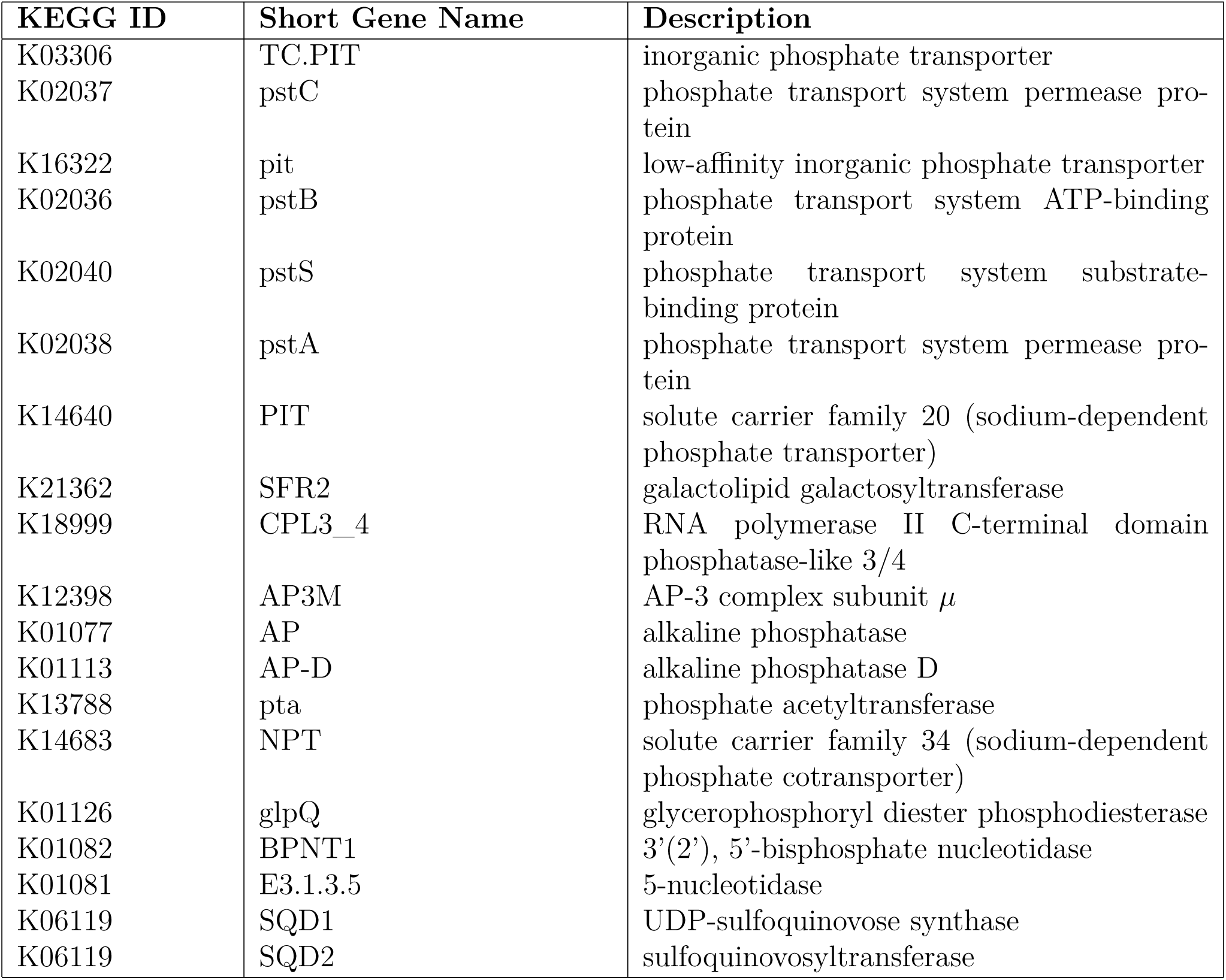
IDs of phosphate uptake and use related genes in the KEGG database.

**Figure 10:**
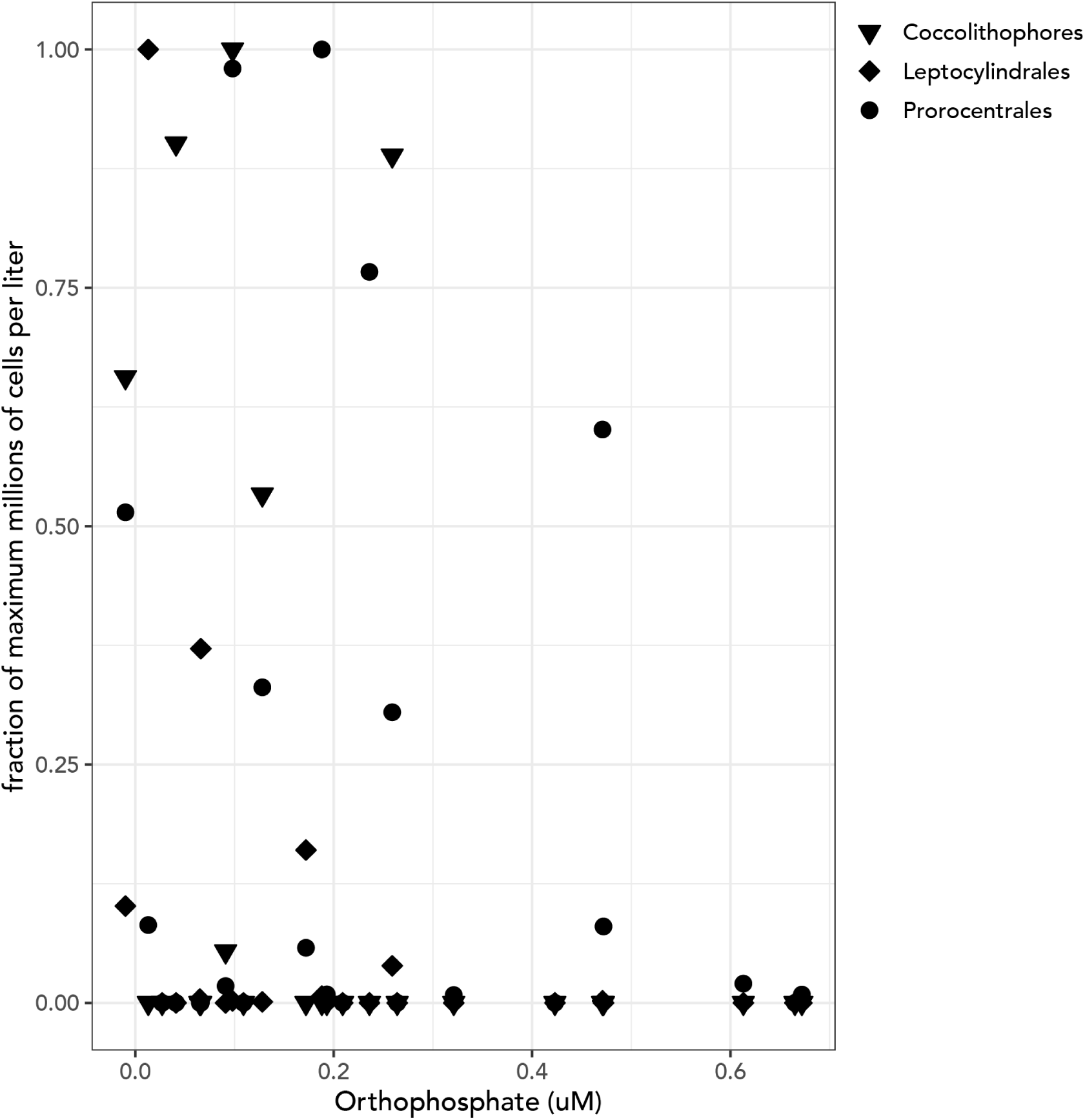
Relationship between scaled relative abundance of coccolithophores, *Leptocylindrales* and *Prorocentrales* (relative counts in millions of cells per milliliter divided by the maximum relative count measured for each group) and the concentration of orthophosphate (*µ*M).

**Table 3:**
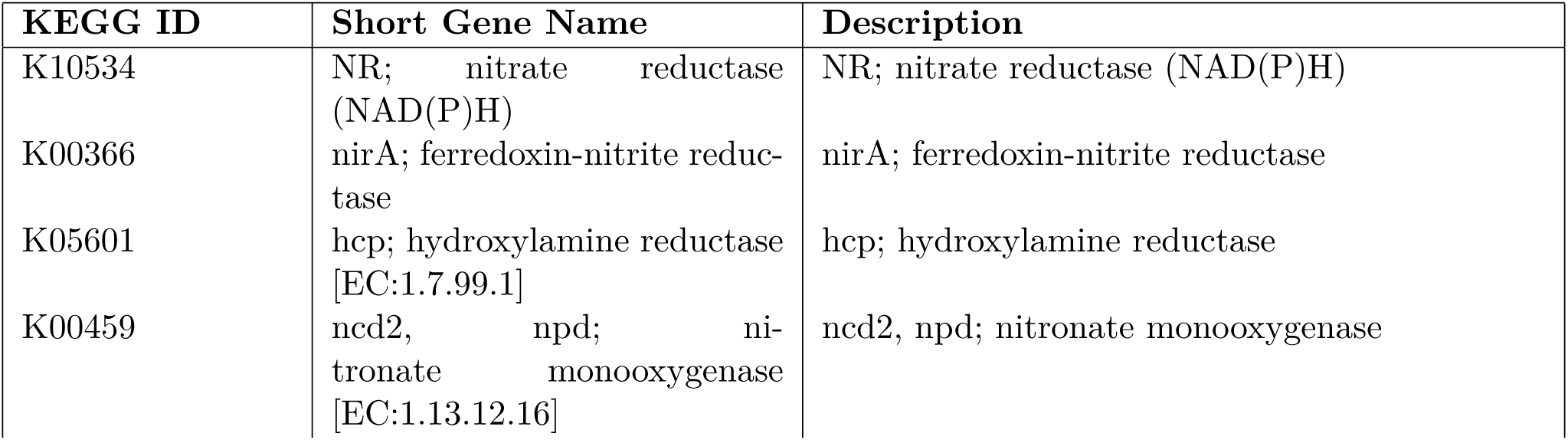
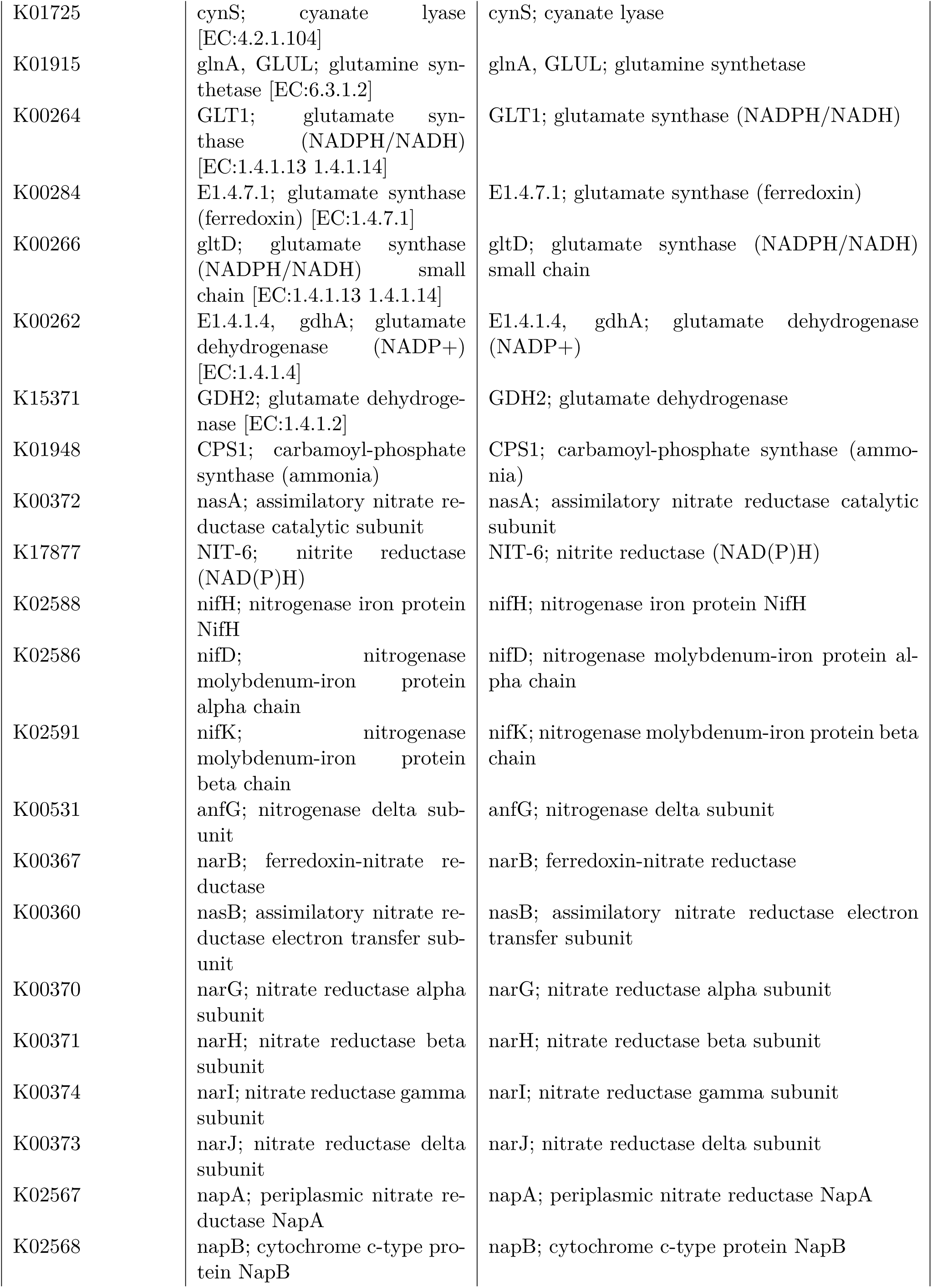
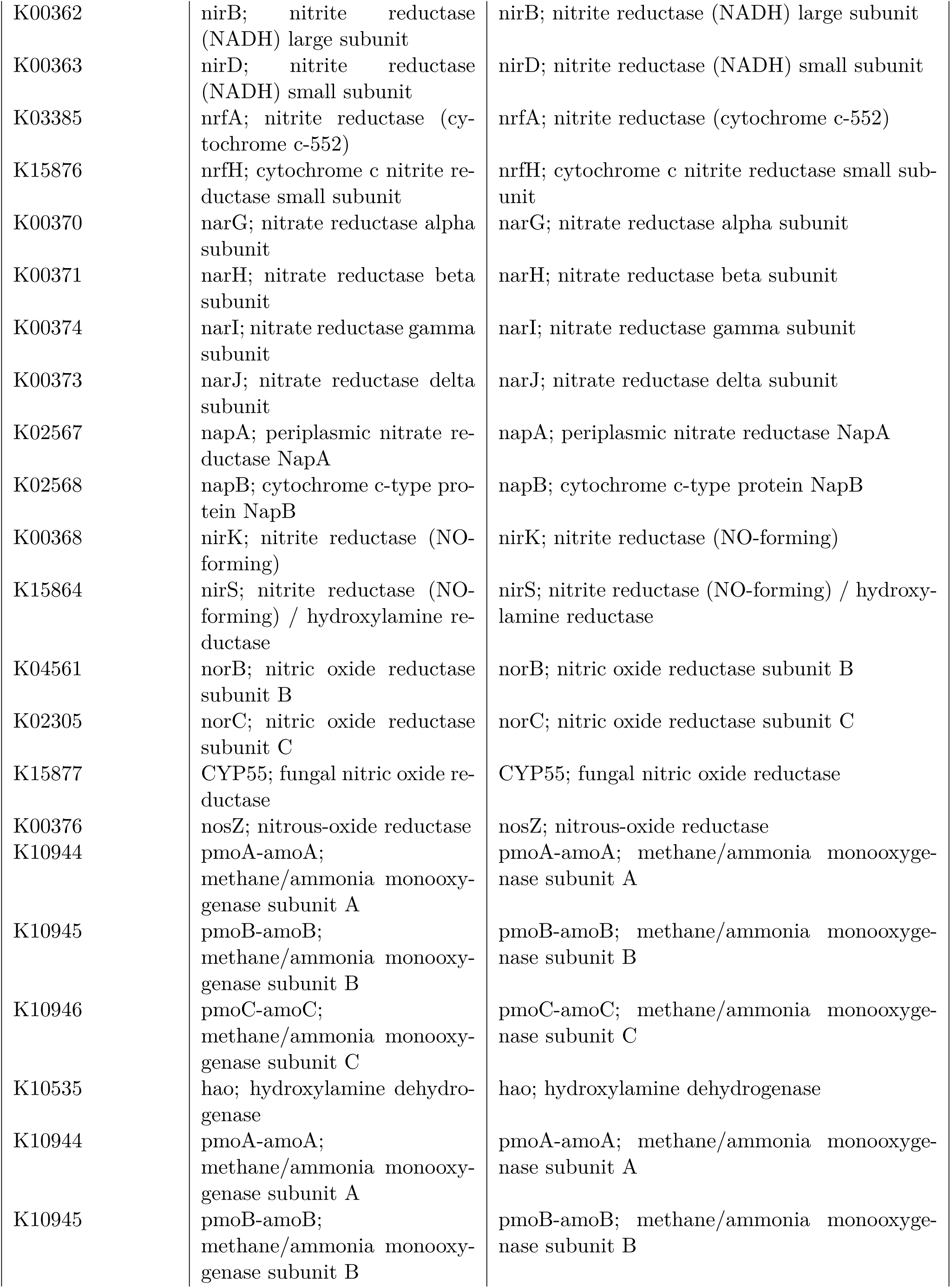
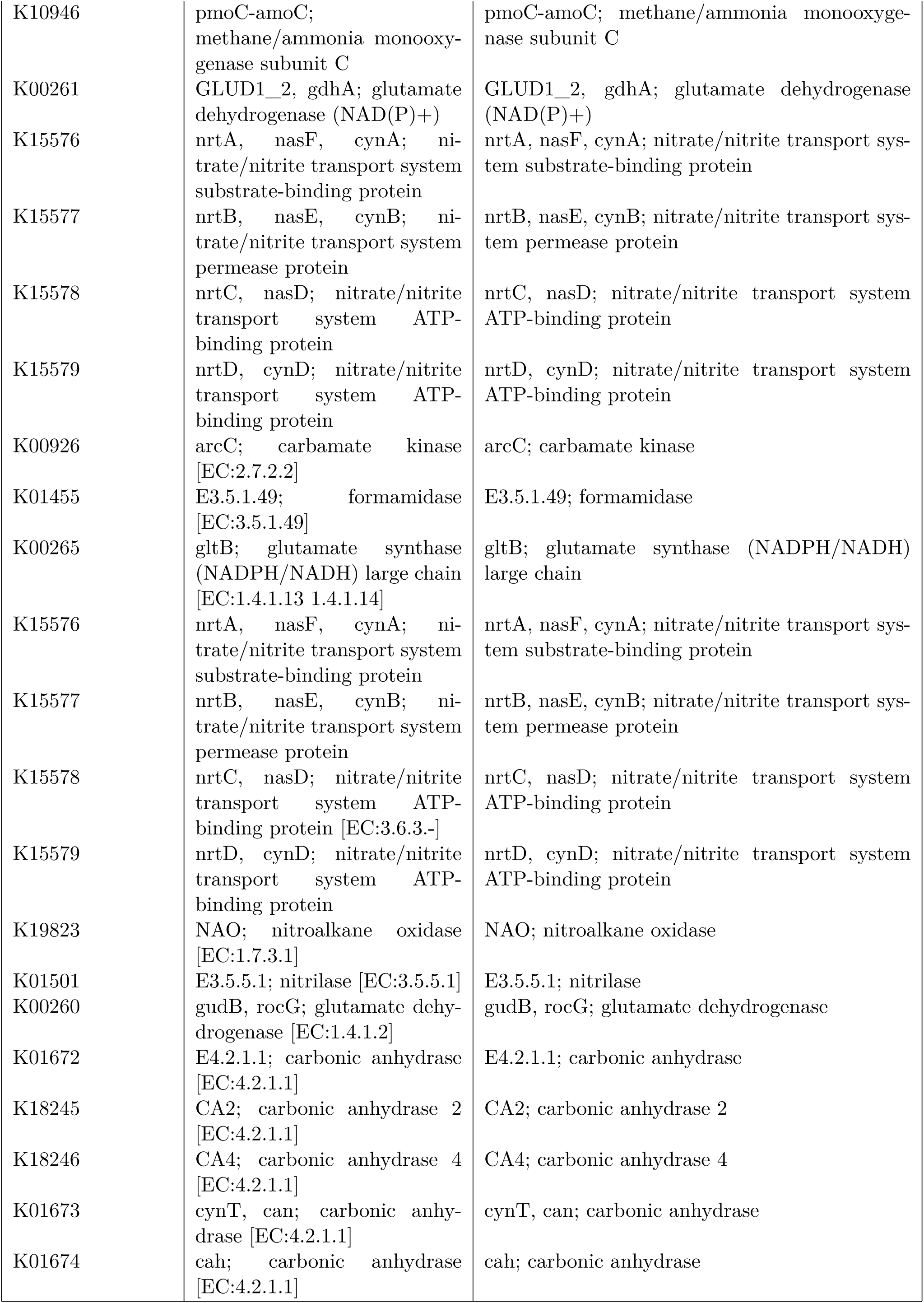
IDs of nitrogen uptake and use related genes in the KEGG database.

**Figure 11:**
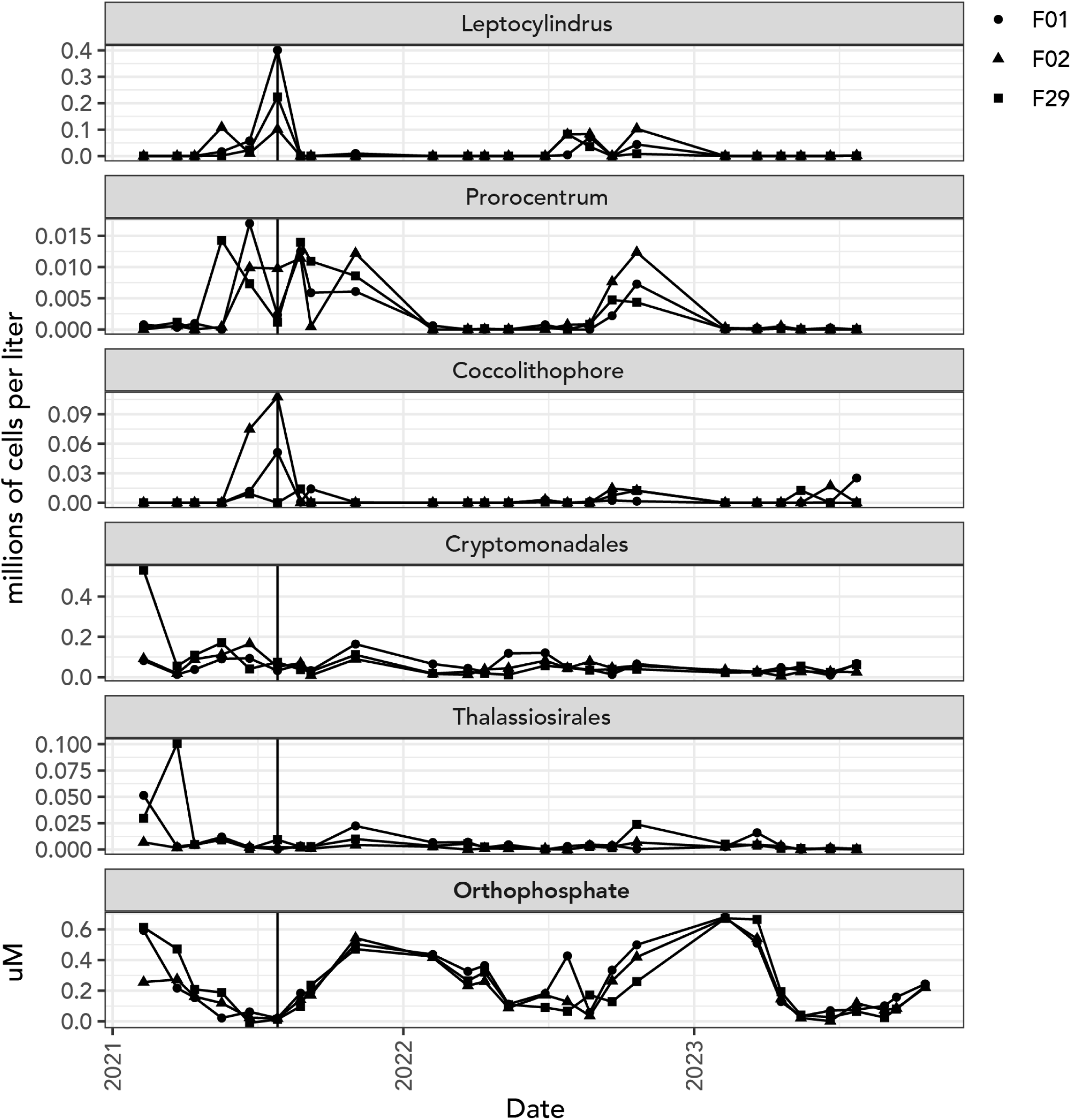
Comparison of microscopic counts for taxa of interest (top 5 panels) as compared to orthophosphate availability (bottom panel). The vertical line shows the July 2021 sampling point. Data for counts ends in July 2023 when metatranscriptomic sampling ended.

#### B.1 Assembly statistics

**Figure 12:**
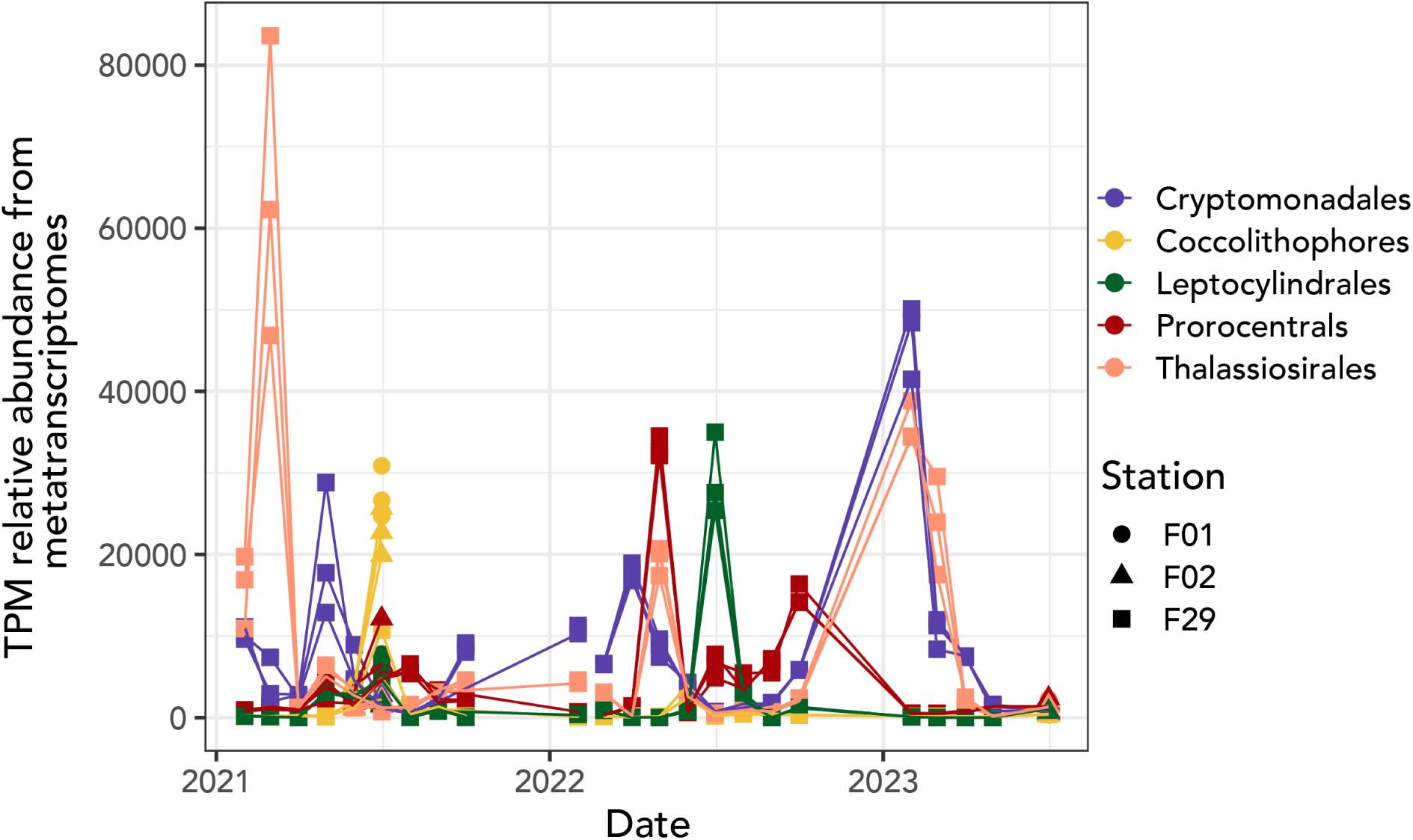
Relative abundance as assessed in transcripts per million (TPM) for the five groups of phytoplankton assessed in the study. This plot highlights the maximum relative gene expression of Thalassiosirales in March of 2021, Prorocentrales in May of 2022, Leptocylindrales in July of 2022, and Cryptomonadales in February of 2023.

**Table 4:** Summary of library size and percent mapping data for the Salmon mapping results for each of the assemblies and replicates from the metatranscriptome study.

| Sample | Month | Year | Station | Replicate | Total library size (in paired reads) | Mapping rate to combined assembly |
| --- | --- | --- | --- | --- | --- | --- |
| S81 | February | 2021 | F29 | 1 | 69430271 | 87.3% |
| S83 | February | 2021 | F29 | 2 | 65049261 | 86.2% |
| S87 | February | 2021 | F29 | 3 | 44854995 | 85.4% |
| S60 | March | 2021 | F29 | 1 | 75276009 | 92.9% |
| S45 | March | 2021 | F29 | 2 | 57237222 | 92.5% |
| S57 | March | 2021 | F29 | 3 | 56927863 | 89.0% |
| S75 | April | 2021 | F29 | 1 | 69765821 | 91.7% |
| S69 | April | 2021 | F29 | 2 | 62818627 | 91.6% |
| S80 | April | 2021 | F29 | 3 | 66769929 | 91.6% |
| S54 | May | 2021 | F29 | 1 | 78987969 | 91.8% |
| S56 | May | 2021 | F29 | 2 | 55435175 | 88.3% |
| S53 | May | 2021 | F29 | 3 | 46787893 | 92.0% |
| S76 | June | 2021 | F02 | 1 | 50475796 | 77.7% |
| S74 | June | 2021 | F02 | 2 | 49760061 | 78.4% |
| S79 | June | 2021 | F02 | 3 | 41087478 | 78.4% |
| S64 | June | 2021 | F29 | 1 | 47914370 | 81.9% |
| S68 | June | 2021 | F29 | 2 | 81150293 | 87.8% |
| S63 | June | 2021 | F29 | 3 | 65281603 | 86.9% |
| S36 | July | 2021 | F01 | 1 | 48654324 | 86.9% |
| S31 | July | 2023 | F01 | 2 | 56129265 | 86.2% |
| S32 | July | 2023 | F01 | 3 | 52039434 | 85.3% |
| S30 | July | 2023 | F02 | 1 | 50400305 | 86.2% |
| S29 | July | 2023 | F02 | 2 | 56662968 | 86.5% |
| S43 | July | 2023 | F02 | 3 | 52404342 | 85.4% |
| S26 | July | 2021 | F29 | 1 | 23535883 | 87.1% |
| S42 | July | 2021 | F29 | 2 | 55937824 | 90.2% |
| S40 | July | 2021 | F29 | 3 | 92204749 | 90.4% |
| S27 | August | 2021 | F29 | 1 | 59356257 | 82.3% |
| S38 | August | 2021 | F29 | 2 | 63738809 | 82.9% |
| S35 | August | 2021 | F29 | 3 | 71178928 | 85.6% |
| S5 | September | 2021 | F29 | 1 | 56064468 | 86.3% |
| S2 | September | 2021 | F29 | 2 | 65515160 | 85.7% |
| S1 | September | 2021 | F29 | 3 | 96974973 | 85.6% |
| S16 | October | 2021 | F29 | 1 | 62830142 | 81.2% |
| S17 | October | 2021 | F29 | 2 | 66078863 | 80.3% |
| S4 | October | 2021 | F29 | 3 | 71130497 | 80.1% |
| S15 | February | 2022 | F29 | 1 | 80492347 | 90.8% |
| S34 | February | 2022 | F29 | 2 | 53366852 | 91.8% |
| S125 | March | 2022 | F29 | 1 | 55564658 | 96.3% |
| S124 | March | 2022 | F29 | 2 | 73850577 | 96.3% |
| S106 | March | 2022 | F29 | 3 | 65716100 | 95.4% |
| S103 | April | 2022 | F29 | 1 | 76199564 | 96.1% |
| S104 | April | 2022 | F29 | 2 | 72896189 | 96.0% |
| S102 | April | 2022 | F29 | 3 | 68616435 | 95.4% |
| S98 | May | 2022 | F29 | 1 | 64617309 | 94.4% |
| S95 | May | 2022 | F29 | 2 | 62681097 | 94.9% |
| S96 | May | 2022 | F29 | 3 | 82026055 | 94.6% |
| S133 | June | 2022 | F29 | 1 | 123957116 | 92.5% |
| S129 | June | 2022 | F29 | 2 | 51829818 | 86.1% |
| S132 | June | 2022 | F29 | 3 | 94521619 | 91.9% |
| S113 | July | 2022 | F29 | 1 | 55869793 | 88.4% |
| S117 | July | 2022 | F29 | 2 | 54251245 | 87.1% |
| S118 | July | 2022 | F29 | 3 | 70315527 | 87.2% |
| S139 | August | 2022 | F29 | 1 | 73309296 | 85.0% |
| S140 | August | 2022 | F29 | 2 | 41182477 | 78.7 % |
| S137 | August | 2022 | F29 | 3 | 73334860 | 84.9% |
| S149 | September | 2022 | F29 | 1 | 45120678 | 88.5% |
| S150 | September | 2022 | F29 | 2 | 54550587 | 89.2% |
| S143 | September | 2022 | F29 | 3 | 49402105 | 84.8% |
| S158 | October | 2022 | F29 | 1 | 44189274 | 77.7% |
| S156 | October | 2022 | F29 | 2 | 41659328 | 77.2% |
| S159 | October | 2022 | F29 | 3 | 46100301 | 76.9% |
| S161 | February | 2023 | F29 | 1 | 44885259 | 90.4% |
| S162 | February | 2023 | F29 | 2 | 30048275 | 91.4% |
| S163 | February | 2023 | F29 | 3 | 47300641 | 91.5% |
| S171 | March | 2023 | F29 | 1 | 43997555 | 87.8% |
| S172 | March | 2023 | F29 | 2 | 47234924 | 87.3% |
| S170 | March | 2023 | F29 | 3 | 41625426 | 87.2% |
| S179 | April | 2023 | F29 | 1 | 59631248 | 93.4% |
| S181 | April | 2023 | F29 | 2 | 46118030 | 93.5% |
| S180 | April | 2023 | F29 | 3 | 47267741 | 97.8% |
| S189 | May | 2023 | F29 | 1 | 23629839 | 95.2% |
| S190 | May | 2023 | F29 | 2 | 25031184 | 95.6% |
| S188 | May | 2023 | F29 | 3 | 19238470 | 96.4% |
| S201 | July | 2023 | F01 | 1 | 55186880 | 88.0% |
| S202 | July | 2023 | F01 | 2 | 63519049 | 87.8% |
| S200 | July | 2023 | F01 | 3 | 52642600 | 88.4% |
| S203 | July | 2023 | F02 | 1 | 47970386 | 87.0% |
| S205 | July | 2023 | F02 | 2 | 59092652 | 87.6% |
| S204 | July | 2023 | F02 | 3 | 87392386 | 86.4% |
| S198 | July | 2023 | F29 | 1 | 63857333 | 87.7% |
| S197 | July | 2023 | F29 | 2 | 54537555 | 88.4% |
| S199 | July | 2023 | F29 | 3 | 66466593 | 87.7% |

## References

Anderson, Donald M (1997). “Bloom dynamics of toxic Alexandrium species in the northeastern US”. In: Limnology and Oceanography 42.5part2, pp. 1009–1022.

Andrews, Simon et al. (Jan. 2012). FastQC. Babraham Institute. Babraham, UK.

Aramaki, Takuya et al. (2020). “KofamKOALA: KEGG Ortholog assignment based on profile HMM and adaptive score threshold”. In: Bioinformatics 36.7, pp. 2251–2252.

Arar, Elizabeth J and Gary B Collins (1997). *Method 445.0: In vitro determination of chlorophyll a and pheophytin a in marine and freshwater algae by fluorescence*. United States Environmental Protection Agency, Office of Research and.

Barnes, Morvan K et al. (2015). “Drivers and effects of Karenia mikimotoi blooms in the western English Channel”. In: Progress in Oceanography 137, pp. 456–469.

Barth, Alex et al. (2020). “Seasonal and interannual variability of phytoplankton abundance and community composition on the Central Coast of California”. In: Marine Ecology Progress Series 637, pp. 29–43.

Boivin-Rioux, Aude et al. (2021). “Predicting the effects of climate change on the occurrence of the toxic dinoflagellate Alexandrium catenella along canada’s east coast”. In: Frontiers in Marine Science 7, p. 608021.

Bolger, Anthony M, Marc Lohse, and Bjoern Usadel (2014). “Trimmomatic: a flexible trimmer for Illumina sequence data”. In: Bioinformatics 30.15, pp. 2114–2120.

Borkman, David G et al. (2016). “Variability of winter-spring bloom Phaeocystis pouchetii abundance in Massachusetts Bay”. In: Estuaries and Coasts 39.4, pp. 1084–1099.

Bushmanova, Elena et al. (2019). “rnaSPAdes: a de novo transcriptome assembler and its application to RNA-Seq data”. In: GigaScience 8.9, giz100.

Camarena-Gómez, Maria T et al. (2018). “Shifts in phytoplankton community structure modify bacterial production, abundance and community composition”. In: Aquatic Microbial Ecology 81.2, pp. 149–170.

Cliff, Alex et al. (2023). “Polyphosphate synthesis is an evolutionarily ancient phosphorus storage strategy in microalgae”. In: Algal Research 73, p. 103161.

Costa, A, E Larson, and K Stamieszkin (2013). “Quality Assurance Project Plan (QAPP) for Water”. In: MWRA Environmental Water Quality Report.

Costa, A et al. (2023). “Water Column Monitoring Results for Cape Cod”. In.

Costa, Amy and Pat Hughes (2012). “How is Our Bay?” In: Provincetown Center for Coastal Studies.

Dai, Yanhui et al. (2023). “Coastal phytoplankton blooms expand and intensify in the 21st century”. In: Nature 615.7951, pp. 280–284.

Faust, Maria A and Rose A Gulledge (2002). “Identifying harmful marine dinoflagellates”. In.

Fernández-Alías, Alfredo et al. (2022). “Nutrient overload promotes the transition from top-down to bottom-up control and triggers dystrophic crises In a Mediterranean coastal lagoon”. In: Science of the Total Environment 846, p. 157388.

Fielding, Samuel R (2013). “Emiliania huxleyi specific growth rate dependence on temperature”. In: Limnology and Oceanography 58.2, pp. 663–666.

Fu, Fei-Xue et al. (2008). “A comparison of future increased CO2 and temperature effects on sympatric Heterosigma akashiwo and Prorocentrum minimum”. In: Harmful Algae 7.1, pp. 76–90.

Glibert, Patricia M et al. (2016). “Pluses and minuses of ammonium and nitrate uptake and assimilation by phytoplankton and implications for productivity and community composition, with emphasis on nitrogen-enriched conditions”. In: Limnology and Oceanography 61.1, pp. 165–197.

Godrijan, Jelena, David Drapeau, and William M Balch (2020). “Mixotrophic uptake of organic compounds by coccolithophores”. In: Limnology and Oceanography 65.6, pp. 1410–1421.

Guangmao, Ding and Zhang Shufeng (2018). “Ecological characteristics and the causes of Karenia mikimotoi bloom in the Sansha Bay in 2012”. In: Acta Oceanol Sin 40.6, pp. 104–112.

Haas, B and A Papanicolaou (2021). *TransDecoder identifies candidate coding regions within transcript sequences*.

Hallegraeff, Gustaaf M (2003). “Harmful algal blooms: a global overview”. In: Manual on Harmful Marine Microalgae 33, pp. 1–22.

Hao, Weihua et al. (1997). “Regulation of AP-3 function by inositides: identification of phosphatidylinositol 3, 4, 5-trisphosphate as a potent ligand”. In: Journal of Biological Chemistry 272.10, pp. 6393–6398.

Härtel, Heiko, Peter Dörmann, and Christoph Benning (2000). “DGD1-independent biosynthesis of extraplastidic galactolipids after phosphate deprivation in Arabidopsis”. In: Proceedings of the National Academy of Sciences 97.19, pp. 10649–10654.

Hu, Chuanmin, Zhongping Lee, and Bryan Franz (2012). “Chlorophyll algorithms for oligotrophic oceans: A novel approach based on three-band reflectance difference”. In: Journal of Geophysical Research: Oceans 117.C1.

Huerta-Cepas, Jaime et al. (2017). “Fast genome-wide functional annotation through orthology assignment by eggNOG-mapper”. In: Molecular Biology and Evolution 34.8, pp. 2115–2122.

Hunt, Carlton D et al. (2010). “Phytoplankton patterns in Massachusetts Bay—1992– 2007”. In: Estuaries and Coasts 33, pp. 448–470.

James, Chase C et al. (2022). “Influence of nutrient supply on plankton microbiome biodiversity and distribution in a coastal upwelling region”. In: Nature Communications 13.1, p. 2448.

Jiang, Mingshun et al. (2014). “Nutrient input and the competition between *Phaeocystis pouchetii* and diatoms in Massachusetts Bay spring bloom”. In: Journal of Marine Systems 134, pp. 29–44.

Johnson, Lisa K, Harriet Alexander, and C Titus Brown (2019). “Re-assembly, quality evaluation, and annotation of 678 microbial eukaryotic reference transcriptomes”. In: Gigascience 8.4, giy158.

Johnson, Matthew D (2015). “Inducible mixotrophy in the dinoflagellate *Prorocentrum minimum*”. In: Journal of Eukaryotic Microbiology 62.4, pp. 431–443.

Keeling, Patrick J et al. (2014). “The Marine Microbial Eukaryote Transcriptome Sequencing Project (MMETSP): illuminating the functional diversity of eukaryotic life in the oceans through transcriptome sequencing”. In: PLoS Biol 12.6, e1001889.

Kelly, Amélie A and Peter Dörmann (2002). “DGD2, an *Arabidopsis* gene encoding a UDP-galactose-dependent digalactosyldiacylglycerol synthase is expressed during growth under phosphate-limiting conditions”. In: Journal of Biological Chemistry 277.2, pp. 1166–1173.

Kelly, Amélie A, John E Froehlich, and Peter Dörmann (2003). “Disruption of the two digalactosyldiacylglycerol synthase genes DGD1 and DGD2 in Arabidopsis reveals the existence of an additional enzyme of galactolipid synthesis”. In: The Plant Cell 15.11, pp. 2694–2706.

Klemetsen, Terje, et al. (Jan. 2018). “The MAR databases: development and implementation of databases specific for marine metagenomics”. en. In: Nucleic Acids Res. 46.D1, pp. D692–D699.

Krinos, A I, et al. (Mar. 2023). “Reverse engineering environmental metatranscriptomes clarifies best practices for eukaryotic assembly”. en. In: BMC Bioinformatics 24.1, p. 74.

Krinos, Arianna I. et al. (2021). “EUKulele: Taxonomic annotation of the unsung eukaryotic microbes”. In: Journal of Open Source Software 6.57, p. 2817. doi: 10 . 21105/joss.02817. url: 10.21105/joss.02817.

Ladd, Tanika M et al. (2018). “Exposure to oil from the 2015 Refugio spill alters the physiology of a common harmful algal bloom species, Pseudo-nitzschia australis, and the ubiquitous coccolithophore, Emiliania huxleyi”. In: Marine Ecology Progress Series 603, pp. 61–78.

Landa, Marine et al. (2016). “Shifts in bacterial community composition associated with increased carbon cycling in a mosaic of phytoplankton blooms”. In: The ISME Journal 10.1, pp. 39–50.

Li, Dinghua et al. (2015). “MEGAHIT: an ultra-fast single-node solution for large and complex metagenomics assembly via succinct de Bruijn graph”. In: Bioinformatics 31.10, pp. 1674–1676.

Li, Xiaodong et al. (2019). “A review of Karenia mikimotoi: Bloom events, physiology, toxicity and toxic mechanism”. In: Harmful Algae 90, p. 101702.

Libby, PS et al. (2006). “Combined work/quality assurance project plan (QAPP) for water column monitoring 2006–2007, tasks 4, 5, 6, 7, 8, 11”. In: MWRA Environmental Water Quality Report 3, p. 119.

Linge Johnsen, Simen Alexander and Jörg Bollmann (2020). “Coccolith mass and morphology of different Emiliania huxleyi morphotypes: A critical examination using Canary Islands material”. In: PLOS One 15.3, e0230569.

Love, Michael I, Wolfgang Huber, and Simon Anders (2014). “Moderated estimation of fold change and dispersion for RNA-seq data with DESeq2”. In: Genome Biology 15.12, p. 550.

Malboobi, Mohammed Ali and Daniel D Lefebvre (1997). “A phosphate-starvation inducible β-glucosidase gene (psr3. 2) isolated from Arabidopsis thaliana is a member of a distinct subfamily of the BGA family”. In: Plant Molecular Biology 34.1, pp. 57– 68.

Marschall, Harold G (1985). “Comparison of phytoplankton concentrations and cell volume measurements from the continental shelf off Cape Cod, Massachusetts, USA”. In: Hydrobiologia 120.2, pp. 171–179.

Martin, Patrick et al. (2014). “Accumulation and enhanced cycling of polyphosphate by Sargasso Sea plankton in response to low phosphorus”. In: Proceedings of the National Academy of Sciences 111.22, pp. 8089–8094.

Matson, Paul G et al. (2019). “Formation, development, and propagation of a rare coastal coccolithophore bloom”. In: Journal of Geophysical Research: Oceans 124.5, pp. 3298–3316.

Mock, Thomas et al. (2016). “Bridging the gap between omics and earth system science to better understand how environmental change impacts marine microbes”. In: Global Change Biology 22.1, pp. 61–75.

Moran, Mary Ann et al. (2013). “Sizing up metatranscriptomics”. In: The ISME journal 7.2, pp. 237–243.

Morey, Jeanine S et al. (2011). “Transcriptomic response of the red tide dinoflagellate, Karenia brevis, to nitrogen and phosphorus depletion and addition”. In: BMC Genomics 12.1, p. 346.

Oczkowski, Autumn et al. (2018). “How the distribution of anthropogenic nitrogen has changed in Narragansett Bay (RI, USA) following major reductions in nutrient loads”. In: Estuaries and Coasts 41.8, pp. 2260–2276.

Paasche, E (1998). “Roles of nitrogen and phosphorus in coccolith formation in *Emiliania huxleyi* (Prymnesiophyceae)”. In: European Journal of Phycology 33.1, pp. 33– 42.

Paasche, E (2001). “A review of the coccolithophorid Emiliania huxleyi (Prymnesiophyceae), with particular reference to growth, coccolith formation, and calcification-photosynthesis interactions”. In: Phycologia 40.6, pp. 503–529.

Patro, Rob et al. (2017). “Salmon provides fast and bias-aware quantification of transcript expression”. In: Nature Methods 14.4, pp. 417–419.

Pershing, Andrew J et al. (2015). “Slow adaptation in the face of rapid warming leads to collapse of the Gulf of Maine cod fishery”. In: Science 350.6262, pp. 809–812.

Pershing, Andrew J et al. (2021). “Climate impacts on the Gulf of Maine ecosystem: a review of observed and expected changes in 2050 from rising temperatures”. In: Elem Sci Anth 9.1, p. 00076.

Prasse, Jennifer et al. (2004). “Combined Work/Quality Assurance Project Plan (CWQAPP)”. In: MWRA Environmental Water Quality Report.

Sah, Saroj Kumar et al. (2024). “Physiological functions of phospholipid: diacylglycerol acyltransferases”. In: Plant and Cell Physiology 65.6, pp. 863–871.

Scully, Malcolm E et al. (2022). “Unprecedented summer hypoxia in southern Cape Cod Bay: an ecological response to regional climate change?” In: Biogeosciences 19.14, pp. 3523–3536.

Shemi, Adva et al. (2016). “Phosphorus starvation induces membrane remodeling and recycling in Emiliania huxleyi”. In: New Phytologist 211.3, pp. 886–898.

Smayda, Theodore J (2011). “Cryptic planktonic diatom challenges phytoplankton ecologists”. In: Proceedings of the National Academy of Sciences 108.11, pp. 4269–4270.

Smith Jr, Walker O et al. (2021). “A regional, early spring bloom of Phaeocystis pouchetii on the New England continental shelf”. In: Journal of Geophysical Research: Oceans 126.2.

Steinegger, Martin and Johannes Söding (2017). “MMseqs2 enables sensitive protein sequence searching for the analysis of massive data sets”. In: Nature Biotechnology 35.11, pp. 1026–1028.

Stoecker, Diane K et al. (2017). “Mixotrophy in the marine plankton”. In: Annual Review of Marine Science 9, pp. 311–335.

Su, Jinzhu et al. (2024). “Environmental drivers and prediction of Karenia mikimotoi proliferation in coastal area, Southeast China”. In: Marine Biology 171.2, p. 57.

Telesh, Irena V, Hendrik Schubert, and Sergei O Skarlato (2016). “Ecological niche partitioning of the invasive dinoflagellate Prorocentrum minimum and its native congeners in the Baltic Sea”. In: Harmful Algae 59, pp. 100–111.

Thompson, PA et al. (2009). “Long-term changes in temperate Australian coastal waters: implications for phytoplankton”. In: Marine Ecology Progress Series 394, pp. 1– 19.

Thorlby, Glenn, Nicolas Fourrier, and Gareth Warren (2004). “The SENSITIVE TO FREEZING2 gene, required for freezing tolerance in Arabidopsis thaliana, encodes a β-glucosidase”. In: The Plant Cell 16.8, pp. 2192–2203.

Townsend, David W (1991). “Influences of oceanographic processes on the biological productivity of the Gulf of Maine.” In: Reviews in Aquatic Sciences 5.3, pp. 211– 230.

Van Mooy, Benjamin AS et al. (2009). “Phytoplankton in the ocean use non-phosphorus lipids in response to phosphorus scarcity”. In: Nature 458.7234, pp. 69–72.

Veldhuis, MJW and W Admiraal (1987). “Influence of phosphate depletion on the growth and colony formation of Phaeocystis pouchetii”. In: Marine Biology 95.1, pp. 47–54.

Veldhuis, MJW, F Colijn, and W Admiraal (1991). “Phosphate utilization in Phaeocystis pouchetii (Haptophyceae)”. In: Marine Ecology 12.1, pp. 53–62.

Yard, Charlestown Navy et al. (2002). “Combined Work/Quality Assurance Project Plan (CWQAPP)”. In: MWRA Environmental Water Quality Report.

Yentsch, Charles S and David W Menzel (1963). “A method for the determination of phytoplankton chlorophyll and phaeophytin by fluorescence”. In: Deep Sea Research and Oceanographic Abstracts. Vol. 10. 3. Elsevier, pp. 221–231.

Zampieri, Guido et al. (2023). “Metatranscriptomics-guided genome-scale metabolic modeling of microbial communities”. In: Cell Reports Methods 3.1.

